# Structural snapshots of La Crosse virus polymerase reveal the mechanisms underlying *Peribunyaviridae* replication and transcription

**DOI:** 10.1101/2021.07.20.453038

**Authors:** Benoît Arragain, Quentin Durieux Trouilleton, Florence Baudin, Stephen Cusack, Guy Schoehn, Hélène Malet

**Affiliations:** Univ. Grenoble Alpes, CNRS, CEA, IBS, F-38000 Grenoble; European Molecular Biology Laboratory (EMBL), Structural and Computational Biology Unit, Heidelberg, Germany; European Molecular Biology Laboratory (EMBL), Grenoble, France; Institut Universitaire de France (IUF).

## Abstract

Segmented negative-strand RNA bunyaviruses encode a multi-functional polymerase that performs genome replication and transcription. Here, we establish conditions for *in vitro* activity of La Crosse virus polymerase and visualize by cryo-electron microscopy its conformational dynamics, unveiling the precise molecular mechanics underlying its essential activities. Replication initiation is coupled to distal duplex promoter formation, endonuclease movement, prime-and-realign loop extension and closure of the polymerase core that direct the template towards the active site. Transcription initiation depends on C-terminal region closure and endonuclease movements that firstly prompt primer cleavage and secondly promote primer entry in the active site. Product realignment after priming, observed in replication and transcription, is triggered by the prime-and-realign loop. Switch to elongation results in polymerase reorganization and core region opening to facilitate template-product duplex formation in the active site cavity. The detailed uncovered mechanics will be crucial for future design of antivirals counteracting bunyaviral life-threatening pathogens.

## INTRODUCTION

*Bunyavirales* is a large order of segmented negative sense single stranded RNA viruses (sNSV). It comprises 12 families and more than 500 viruses, amongst which human dangerous pathogens such as La Crosse (LACV, *Peribunyaviridae* family), Hantaan (HTNV, *Hantaviridae* family), Rift Valley Fever, Severe Fever with Thrombocytopenia Syndrome (RVFV and SFTSV, *Phenuiviridae* family), Lassa (LASV, *Arenaviridae* family) and Crimean Congo Haemorrhagic fever viruses (CCHFV, *Nairoviridae* family)^1,2^. There is currently no treatment or vaccine to counteract them. In this context, we focus our interest on LACV and more precisely on essential steps of its viral cycle: genome replication and transcription. These processes are catalysed by the virally-encoded RNA-dependent RNA-polymerase (RdRp), also called L protein (LACV-L)^3^. This large 260kDa multi-functional and monomeric enzyme performs replication of the viral genome (vRNA) into a complementary RNA (cRNA), which is then used as a template to generate nascent vRNA that have a size and composition identical to the parental vRNA. Replication initiation is performed *de novo*, in the absence of primer. LACV-L and other *Peribunyaviridae* are suspected to initiate their replication internally at position 4 of the RNA template to produce a primer that then realigns to the template extremity. This process, called “prime-and-realign”, is made possible by a triplet nucleotide repetition at the 3’-vRNA template extremity (3’-UCAUCA…-5’ for LACV) and has been reported for several families in the *Bunyavirales* order, although with family-dependent specificities^4–6^. Concerning transcription, LACV-L catalyses its initiation through a cap-snatching mechanism, whereby LACV-L steals cellular capped mRNA, binds them through its cap-binding domain (CBD), cleaves them 9 to 17 nucleotides downstream the cap with its endonuclease domain (ENDO), before using them as a primer for transcription initiation^7,8^. The prime-and-realign mechanism has also been observed at this stage resulting in the insertion of a triplet (5’-AGU-3’) or multiple triplets (5’-AGUAGU…-3’) complementary to the 3’-template inserted between the cap-snatched mRNA primer and the viral RNA transcript^7,9,10^.

Structural determination of these essential and complex sNSV polymerases has long been a challenge in the field. Following determination of the X-ray structure of the C-terminally truncated construct of LACV-L^11^, structures of full-length LACV-L, LASV-L, Machupo-L (MACV-L) and SFTSV-L have been determined by cryo-EM^12–15^. They contain a conserved central core comprising the polymerase active site, an N-terminal ENDO and a flexible C-terminal region (CTER) that includes the CBD. They were solved in their apo (SFTSV-L) or pre-initiation form (LACV-L, Machupo-L and LASV-L) with the 3’-vRNA promoters bound in a secondary site at the surface of the polymerases, away from the RNA synthesis active site. The 5’-vRNA promoter, present only in LACV-L structures, is bound as a hook in a distinct and specific site on the polymerase surface.

These structures were important advances in the field, but they do not shed light on the detailed mechanisms by which this key multi-functional molecular machine performs genome replication and transcription. In the present article, we uncover the conditions necessary for LACV-L *in vitro* activity and determine LACV-L structures stalled at specific stages of both replication and transcription revealing with unprecedented detail the critical elements triggering activation of these essential processes.

## RESULTS

### LACV-L replication activity

Visualization of LACV-L activity necessitated careful optimization of (i) the construct, (ii) the tag position and (iii) the composition of the vRNA promoters. Using a mini-replicon system, LACV-L constructs with an N- or C-terminal tag were found to display low activity compared to the construct without tag (**Fig. 1a**). Addition of a Strep-tag in an internal exposed loop called the California insertion (LACV-L_CItag_) displayed 85% activity compared to the LACV-L without tag and was thus chosen for subsequent purification and *in vitro* activity assays (**Fig. 1a**). To abolish *in vitro* unspecific RNA degradation, we introduced the H34K mutation known to prevent ENDO activity^16^. Despite the construct optimization, the homogeneously purified LACV-L_CItag_H34K_ (**Supplementary Fig. 1a**) incubated with wild-type (WT) 5’/3’-vRNA promoters, NTPs and MgCl_2_ at 30° for 4h did not generate replication products (**Fig. 1b, lane 1)**. This behaviour can be explained by the high complementarity of the 5’/3’-vRNA promoters that tend to form stable double-stranded RNA, preventing their binding as single-stranded RNA in their specific binding sites on LACV-L. This situation is specific to *in vitro* reconstitution as *in vivo* each promoter is kept in separate binding sites on the polymerase. To avoid 5’/3’-vRNA duplex formation, we carefully analysed the structure of the 5’-vRNA promoter and its interaction with the polymerase. We mutated the nucleotides G2, U3, A9 and C10 of the 5’-end into C2, G3, C9 and G10, thereby preserving the hook structure and its interaction with LACV-L while significantly decreasing the 5’/3’-vRNA complementarity (**Fig. 1c,** pre-initiation vs initiation). Equivalent activity assays with a 17-base-pair mutated 5’ (5’-1-17BPm) in place of the WT 5’-vRNA led to the formation of two main replication products: one with the expected length of 25 nucleotides, that corresponds to the size of the 3’-vRNA template, and one with a 28-nucleotide length (**Fig. 1b, lane 2)**. Although an extension by three nucleotides is not expected *in vivo*, this result further suggests a prime-and-realign mechanism during replication initiation. Visualisation of clear replication activity with LACV-L_CItag_H34K,_ 5’-1-17BPm and 3’-vRNA1-25 led us to optimize the assay revealing a maximal activity with 2 to 5 mM Mg^2+^ while more abortive products and fewer 25-mer replication products are visualized in presence of Mn^2+^ (**Supplementary Fig. 1b**). A time course indicates that the reaction is optimal at 4h (**Supplementary Fig. 1c**). We then performed equivalent reaction assays omitting either the 5’- or the 3’-vRNA. These resulted in the absence of 25-mer product formation, clearly indicating the requirement of both promoter ends for replication activity (**Fig. 1b, lanes 3, 4**).

**Fig. 1.**
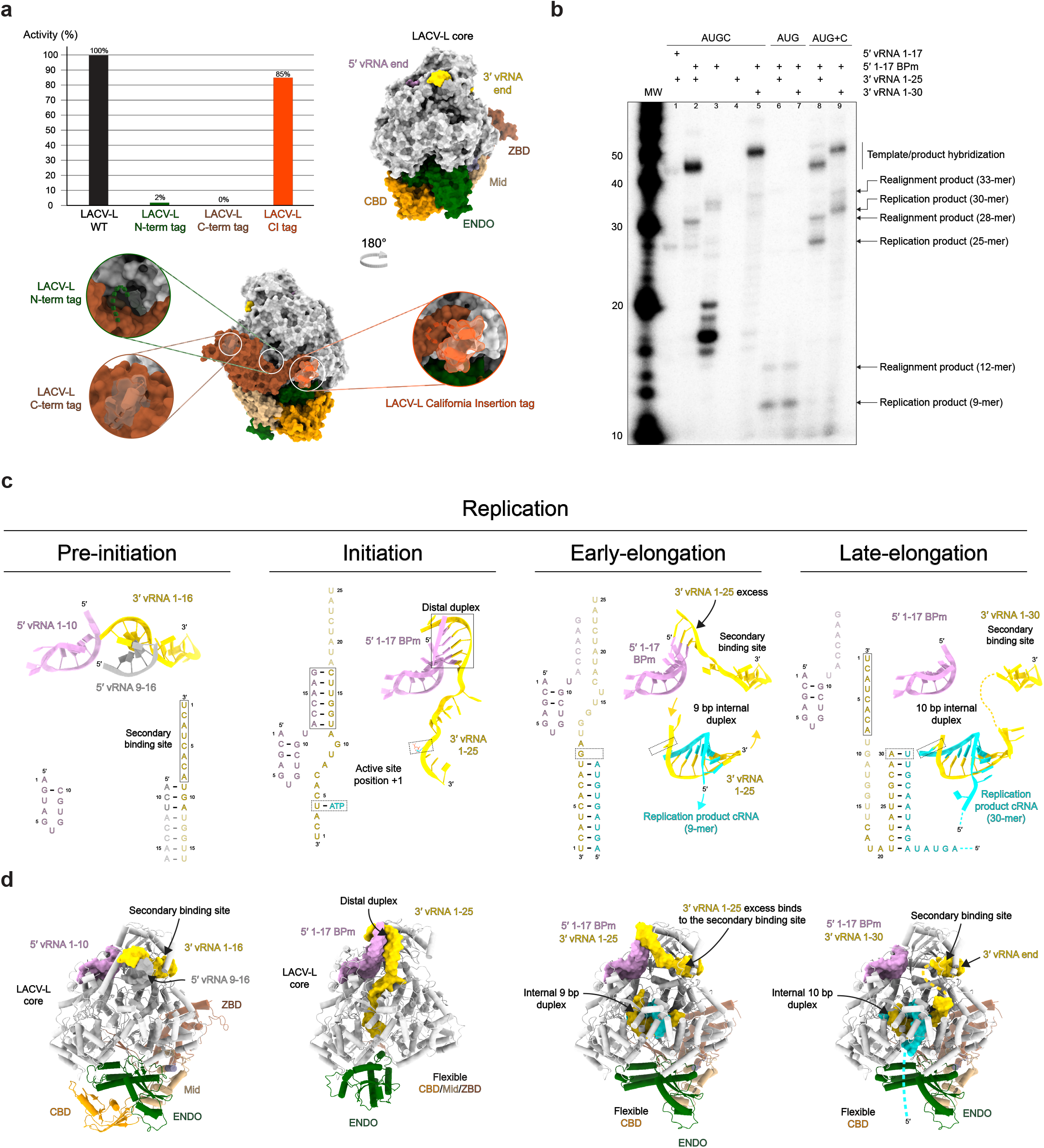
Overview of LACV-L activity and cryo-EM structures stalled in specific replication states. **a,** Histogram of LACV-L mini-replicon activity. The percentage of activity in presence of different tags is compared to LACV-L activity without tag (WT). Tag positions are shown on LACV-L_Nter-Histag_ surface (PDB: 6Z6G) as dotted lines. The polymerase core is colored in light grey with the California insertion in orange, the CBD in gold, the ENDO in green, the mid domain in beige and the ZBD in brown. **b,** *In vitro* replication activity of LACV-L_CItag_H34K_ using different combinations of 5’- and 3’-vRNAs. The assays are done in presence of either the 4 NTPs (AUGC), 3 NTPs (AUG) or 3 NTPs during 4h, subsequently supplemented with CTP for 30 min (AUG+C). Replication products and their respective lengths are displayed on the right side of the gel. The decade molecular weight marker (MW) is shown. **c,** Sequence and secondary structures of the RNA bound to LACV-L_CItag_H34K_ for each replication state. 5’-vRNAs, 3’-vRNAs and replication products are respectively colored in pink/grey, gold and cyan. Bases present in the sequences but not seen in the structures are shown in transparent. **d,** Cartoon representation of each LACV-L_CItag_H34K_ replication structure. LACV-L domains are colored as in **(a)**. RNAs displayed as surfaces are colored as in **(a,c)**. The pre-initiation state corresponds to the PDB 6Z6G.

The dependence of the product size on the template size was analysed, and reveals that, as expected, a 3’-vRNA1-30 template gives rise to products 5 nucleotides longer than a 3’-vRNA1-25 template (**Fig. 1b, lanes 2, 5)**. Incubation of LACV-L_CItag_H34K_ with 5’-1-17BPm, 3’vRNA1-25, ATP, UTP and GTP generates a 9-mer product, consistently with the template sequence that requires CTP incorporation at position 10 (**Fig. 1b, lanes 6, 7)**. Interestingly, CTP addition after 1h restores complete product formation, indicating that the reaction performed with ATP, UTP and GTP is stalled in a physiological, early-elongation state (**Fig. 1b, lanes 8, 9)**. Higher molecular weight bands appear in addition to the expected products (**Fig. 1b, lanes 2, 5, 8, 9)** and likely correspond to the template/product hybridization upon assay arrest due to their high complementary, as reported in several sNSV *in vitro* activity assays ^14,17^.

### Overview of the actively replicating LACV-L structures

Visualisation of replication activity prompted us to collect three large cryo-EM datasets which, coupled with advanced image processing involving extensive 3D classifications, captured LACV-L structures in three key states of replication: initiation, early-elongation and late-elongation at respectively 2.8, 2.9 and 3.9 Å resolution (**Fig. 1c, d, Supplementary Table 1, Supplementary Movie 1**). To stabilize the replication initiation state, LACV-L_CItag_H34K_ was incubated with the 5’-1-17BPm and the 3’-vRNA1-25 in presence of the initial nucleotides to be incorporated. An initiation-mimicking state, with ATP present at position +1 of the active site, was obtained when the LACV-L initiation complex was formed in presence of ATP and UTP (**Fig. 1c, d, Supplementary Fig. 2**). For the replication early-elongation state, LACV-L_CItag_H34K_ was incubated with the previously used RNAs supplemented with ATP, UTP and GTP resulting in the formation of a 9-base pair elongating template-product duplex in the polymerase internal cavity (**Fig. 1c, d, Supplementary Fig. 3**). For the late-elongation state, a 5’-1-17BPm, 3’-vRNA-1-30 and four nucleotides were used. In this state, the 3’-vRNA is entirely replicated, a 10-base pair template-product duplex is visible in the active site and the 3’-vRNA extremity, following its exit from the active site as a single stranded RNA, binds in the polymerase 3’ extremity secondary binding site (**Fig. 1c, d, Supplementary Fig. 3**).

Comparison of the three structures reveals major conformational changes of LACV-L at initiation compared to its stable conformation acquired at pre-initiation and early-/late-elongation states (**Fig. 1d**). A major movement of the ENDO is visible at initiation, coordinated with the movement of the entire CTER including the CBD that becomes flexible at this stage. Whereas the ENDO, mid and zinc-binding domain (ZBD) gain back their pre-initiation position at elongation, the CBD remains flexible.

### The replication initiation state

The principal novelty in the obtained structures concerns the initiation conformation that provides molecular insights into the mechanics underlying this critical step. The distal duplex, formed between nucleotides 12 to 17 of the 3’/5’-vRNA extremities, plays a critical role in triggering the template positioning in the RNA tunnel entrance (**Fig. 2a**). The distal duplex is held in place by the 5’-vRNA hook (nucleotides 1-10), the vRNA Binding Lobe domain (vRBL), the clamp, the arch and the α-ribbon. The RNA entrance tunnel is surrounded by the vRBL, the fingers, the thumb ring and the bridge domains. Nucleotides G9 to A11 of the template are stabilized in the tunnel by multiple hydrophobic and polar interactions while nucleotides A6, C7 and A8 are less coordinated and adopt alternative conformations. Their visualized mobility might provide the template with the necessary flexibility for its realignment after priming (**Fig. 2b**). The 3’ extremity nucleotides U1 to C5 display clearer density correlated with their higher degree of stabilization by the canonical polymerase motifs (**Fig. 2c**). The 3’ extremity itself interacts by an unexpected but crucial element: the loop 982-995, that is localized between the fingers and the palm domain, and which, unexpectedly, plays the role of a replication priming loop (**Fig. 2c**). Its dual role in priming and realignment (see below) leads us to name this element the prime-and-realign loop (PR loop).

**Fig. 2.**
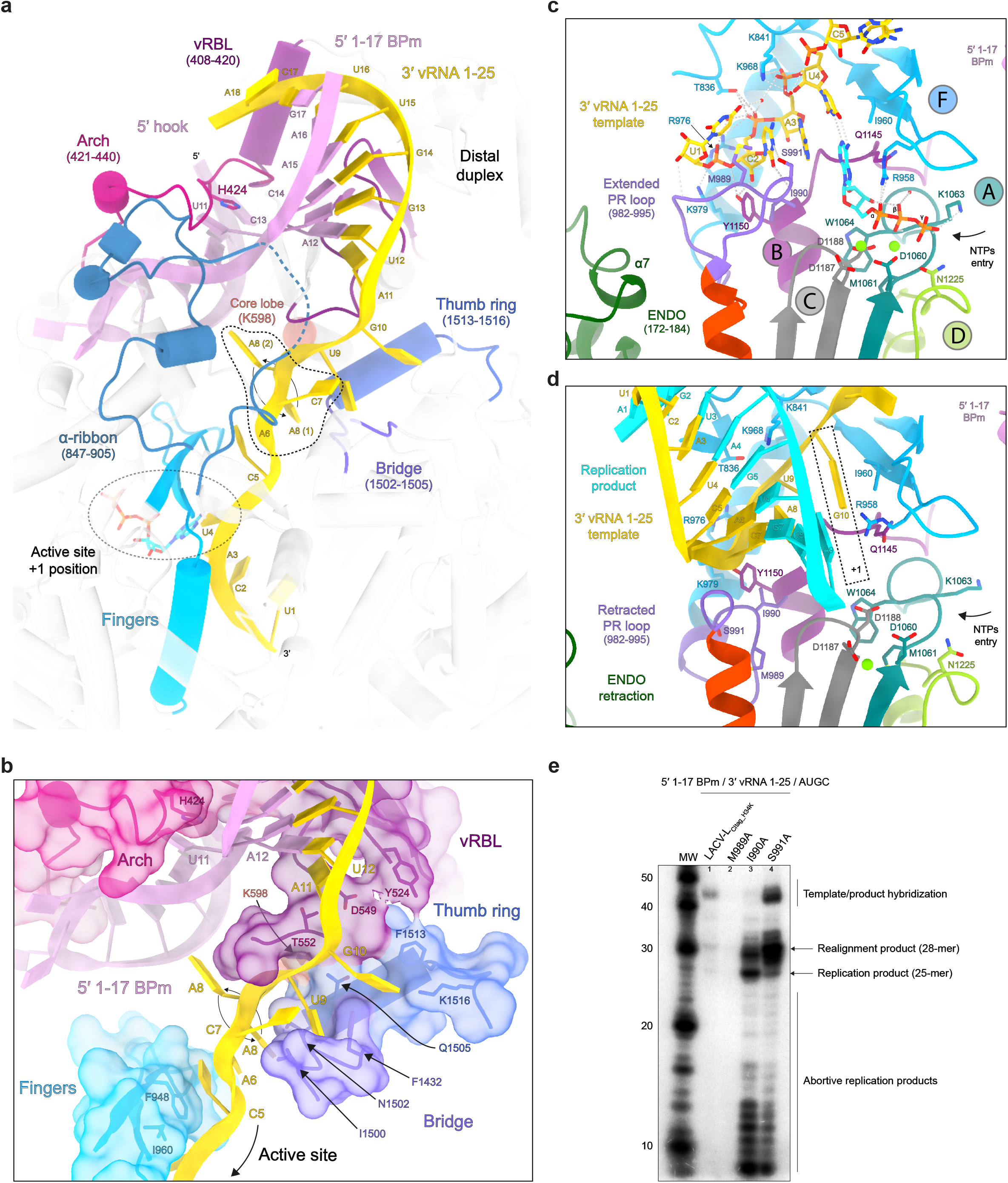
Interactions between LACV-L_CItag_H34K_ and 5’/ 3’-vRNA during replication. **a,** Interaction between LACV-L_CItag_H34K_ and the distal duplex formed by 5’-1-17BPm and 3’-vRNA1-25. Interacting domains, namely the arch, the vRBL, the α-ribbon, the fingers, the bridge and the thumb ring domains are respectively colored in magenta, violet red, steel blue, cyan, purple and blue. The RNAs are colored as in **Fig. 1c,d**. ATP in position +1 of the active site is surrounded by a dotted line. **b,** Zoom on the interaction between LACV-L_CItag_H34K_ and the 3’-vRNA1-25. Surface of interacting domains are displayed and colored according to **(a)**. Flexible nucleotides A6/C7/A8 of the 3’-vRNA1-25 are surrounded by a dotted line. The two visible conformations of A8 are shown. **c,** Active site of LACV-L_CItag_H34K_ in replication initiation state. RdRp motifs A, B, C, D and F are respectively colored in dark turquoise, purple, grey, light green and blue. The extended PR loop and ENDO are colored in orchid and green. Residues that interact with the 3’-vRNA1-25 are displayed. The two magnesium ions are colored in light green. **d,** Active site of LACV-L_CItag_H34K_ in replication elongation state. LACV-L domains, RdRp motifs and vRNAs are colored as in **(c)**. The newly synthesized product forming a 9 base-pair duplex with the 3’-vRNA1-25 template is colored in cyan. The +1 active site position is surrounded by a dotted line. **e,** *In vitro* replication activity of LACV-L_CItag-H34K_, LACV-L_CItag-H34K-M989A_ (M989A), LACV-L_CItag-H34K-I990A_ (I990A), LACV-L_CItag-H34K-S991A_ (S991A) with 5’-1-17BPm, 3’-vRNA1-25 and 4 NTPs. Replication products and their respective lengths are displayed on the right side of the gel. The decade MW marker is shown.

At initiation, the ENDO conformational change brings its residues 172 to 184 (α-helix 7) proximally to the active site triggering the PR loop extension (**Fig. 2c**). The PR loop residue M989 interacts with the 3’ extreme nucleotide U1 in position −3 of the active site while the residues I990 and S991 stabilize the nucleotide C2 in position −2 of the active site (**Fig. 2c**). As the result, the template nucleotides A3 and U4 are localized in position −1 and +1 of the active site (**Figs. 1c, 2c**). The depicted map displays a clear density for an ATP in position +1 that makes hydrogen bonds with U4 and stacks with A3. Its ribose interacts with W1064 (from motif A) and Q1145 (from motif B), its phosphates interact with D1060, M1061, K1063 (from motif A), D1187 (from motif C) and two magnesium ions, themselves located in canonical positions for catalytic magnesium A and B (**Fig. 2c**).

The identification of the unexpected PR loop led us to analyze further its role in replication by engineering single-alanine substitutions of its tip residues M989, I990 and S991 (**Fig. 2e**). M989A abolishes the replication activity, confirming the importance of this residue in precise template positioning (**Fig. 2e, lane 2**). Intriguingly, LACV-L_CItag_H34K _I990A_ is more active than LACV-L_CItag_H34K_ **(Fig. 2e, lane 3 vs lane 1)**. Due to the proximity of I990 to motif B that is implicated in translocation (**Fig. 2c**), it is tempting to speculate that the I990A mutation, by providing more space for the product, may facilitate elongation. LACV-L_CItag_H34K_S991A_, although generating replication product of correct size, 25-mer, leads to more product that are +2, +3 or +4 nucleotides longer **(Fig. 2e, lane 4 vs lane 1**). As S991 interacts in the initiation-mimicking structure with the C2 of the 3’-vRNA, one may suggest the importance of the side chain for proper positioning of the template (**Fig. 2c**). Altogether, these mutations clearly confirm the importance of the PR loop tip in the replication mechanism.

### Switch to elongation and related conformational changes

Progression towards elongation implies important remodeling of LACV-L domains. Retraction of the PR loop, that is coupled with ENDO repositioning (**Fig. 2d, 3a**), leads to the initial formation of the template-product duplex in the active site cavity. The absence of CTP in the mix results in the stalling of LACV-L at template G10 in position +1 of the active site in a post-translocation pre-incorporation state (**Fig. 2d**). Whereas the replication initiation was performed internally, realignment must have occurred as the replication elongation displays an entire product with the 5’-cRNA extremity corresponding to nt 1.

To accommodate the internal 9-base pair duplex, switch from initiation to elongation state results in the opening of the lid, the thumb and the thumb-ring coupled with the extension of the bridge loop 1423-1441 by unwinding of the helix 1435-1438 (**Fig. 3b**). Together these movements switch the core from a closed to an open state, resulting in the extrusion of the previously defined “priming loop” that induces the opening of the template exit tunnel (**Fig. 3c**). As the “priming loop” remains away from the active site during replication and does not appear to be involved in priming in the visualized states, it is rebaptised “template exit plug” (**Fig. 3b, c**). Interestingly, reverse movement from an open to a closed core is observed between pre-initiation and initiation (**Fig. 3b, c**) suggesting that the core closure is a rather transient state acquired at initiation only, while the open core is a more stable long-standing state.

**Fig. 3.**
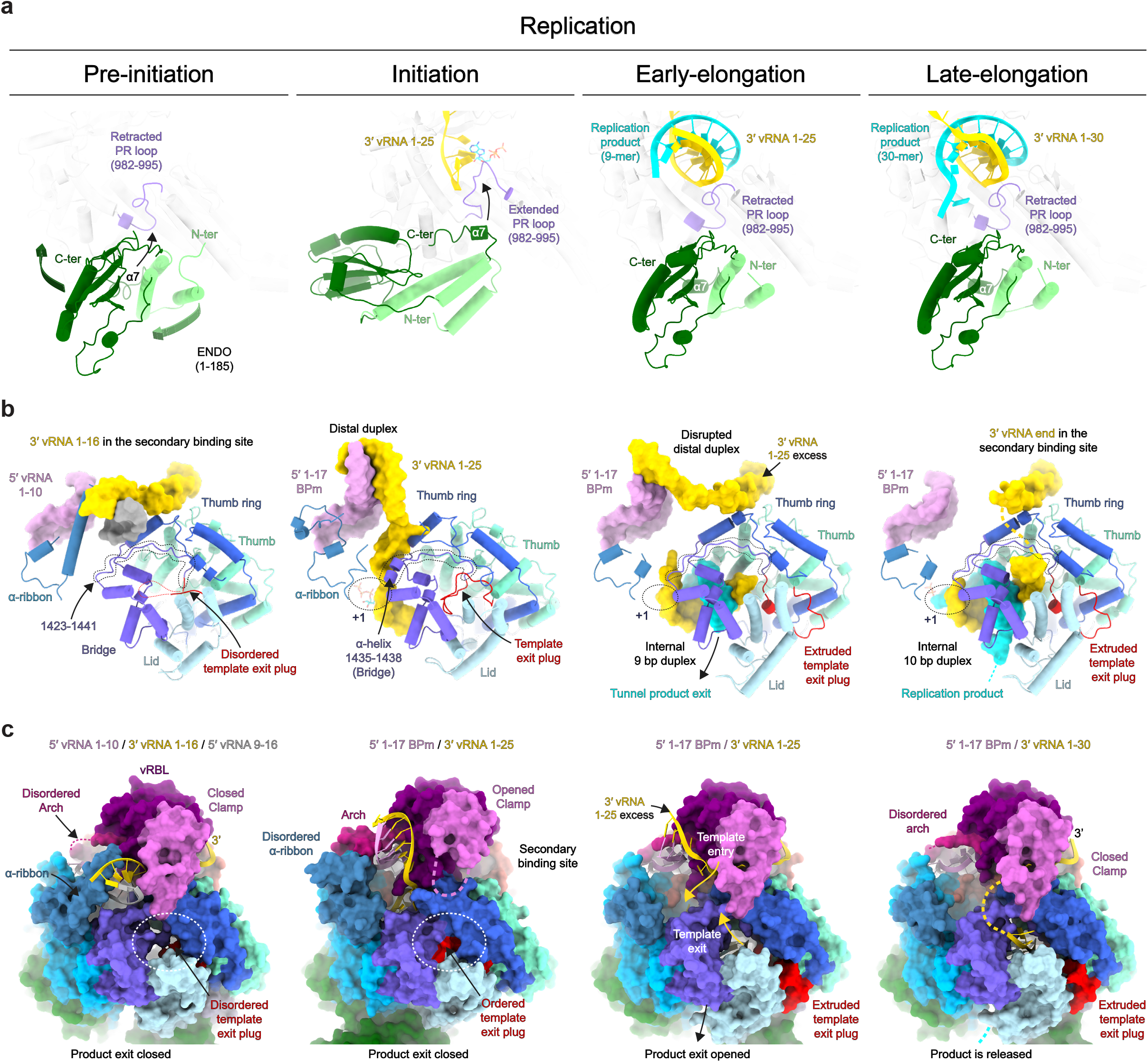
Replication-induced conformational changes of LACV-L_CItag_H34K_. **a,** ENDO and PR loop movements between the replication states. ENDO N-ter and C-ter are indicated. Arrows indicate movements of the ENDO, the α7 helix and the PR loop. **b,** LACV-L domain movements and RNA position variations between the replication states. For clarity, only domains that undergo conformational changes are displayed. The template exit plug, colored in red, is indicated with a dotted line if flexible. RNAs are shown as surfaces and colored as in **Fig. 1c,d**. Bridge residues 1423-1441 are surrounded by a dotted line. **c,** Template exit plug, vRBL, clamp and α-ribbon conformational changes between the replication states. LACV-L surface is displayed and each domain are colored as in **Fig. 2a,b** and **3b**. The template exit tunnel is surrounded by a dotted line. Template entry/exit tunnels are indicated in the early-elongation state with gold arrows.

In addition to the core opening/closing between the replication states, global movements of the vRBL, clamp, arch and α-ribbon are visualised (**Fig. 3b, c**). Ordering of the α-ribbon and closing of the vRBL towards the core at pre-initiation triggers the positioning of the 3’-vRNA in the secondary binding site. At initiation, the acquired α-ribbon flexibility and displacement of the vRBL away from the core results in the disruption of the 3’-vRNA binding in the secondary site. These, coupled with the movements described above, result in the entry of the 3’-vRNA end in the template entry tunnel necessary for initiation. Exit of the 3’ template from the template exit channel at late-elongation triggers a repositioning of the vRBL and the clamp close to the core resulting in the formation of the 3’ secondary binding site as in pre-initiation. The replicated 3’-vRNA extremity exits as a single stranded RNA from the active site cavity, binds to a positively charged groove (**Supplementary Fig. 4**), before reaching the 3’ secondary binding site, ready for the next replication cycle.

### LACV-L transcription activity

Transcription activity was analysed using LACV-L_CItag_H34K_ incubated with a 14-mer capped RNA primer, the 3’-vRNA1-25/1-30 and the 5’-1-17BPm. The chosen capped RNA have a two-(cap14AG) or three-nucleotide (cap14AGU) complementarity with the 3’-vRNA extremity (5’-…ACUACU-3’) to facilitate priming. Time, divalent ion nature and concentration were tested and indicate that the transcription reaction is optimal after 30 min in presence of 2 mM MgCl_2_ (**Supplementary Fig. 5**). Using these conditions, the reactions with cap14AG/cap14AGU lead to the formation of two main transcription products of 37-/36-mer and 40-/39-mer (**Fig. 4a, b, lanes 1, 2**). The minority 37-/36-mer products correspond to the capped RNA size elongated by 23/22 nucleotides, considering the hybridization of the template last 2/3 nucleotides with the cap14AG/capAGU. The majority 40-/39-mer products likely correspond to primed and subsequently realigned transcripts, that result in the addition of three nucleotides at the end of the capped primer before elongation. This is in accordance with the sequencing of capped RNA produced by *Peribunyaviridae* polymerase^7,10,18^. As expected, the size of the transcribed RNA depends on the length of the template: replacing the 3’-vRNA1-25 by a 3’-vRNA1-30 gives rise to products that are 5 nucleotides longer (**Fig. 4a, b, lanes 3, 4**). Usage of a NTP subset containing ATP, UTP and GTP, to obtain an early-stalled transcription complex, shows formation of the expected 21/24-nucleotide products with cap14AG and 20/23 nucleotide products with cap14AGU (**Fig. 4a, b, lanes 5-8**). Interestingly, whereas the realignment was almost complete in presence of the 4 NTP, it is only partial in presence of the 3-NTP subset. Subsequent incubation with a CTP restores formation of full-length capped RNA product, but with a majority of non-realigned capped RNA products (**Fig. 4a, b, lanes 9-12**).

**Fig. 4.**
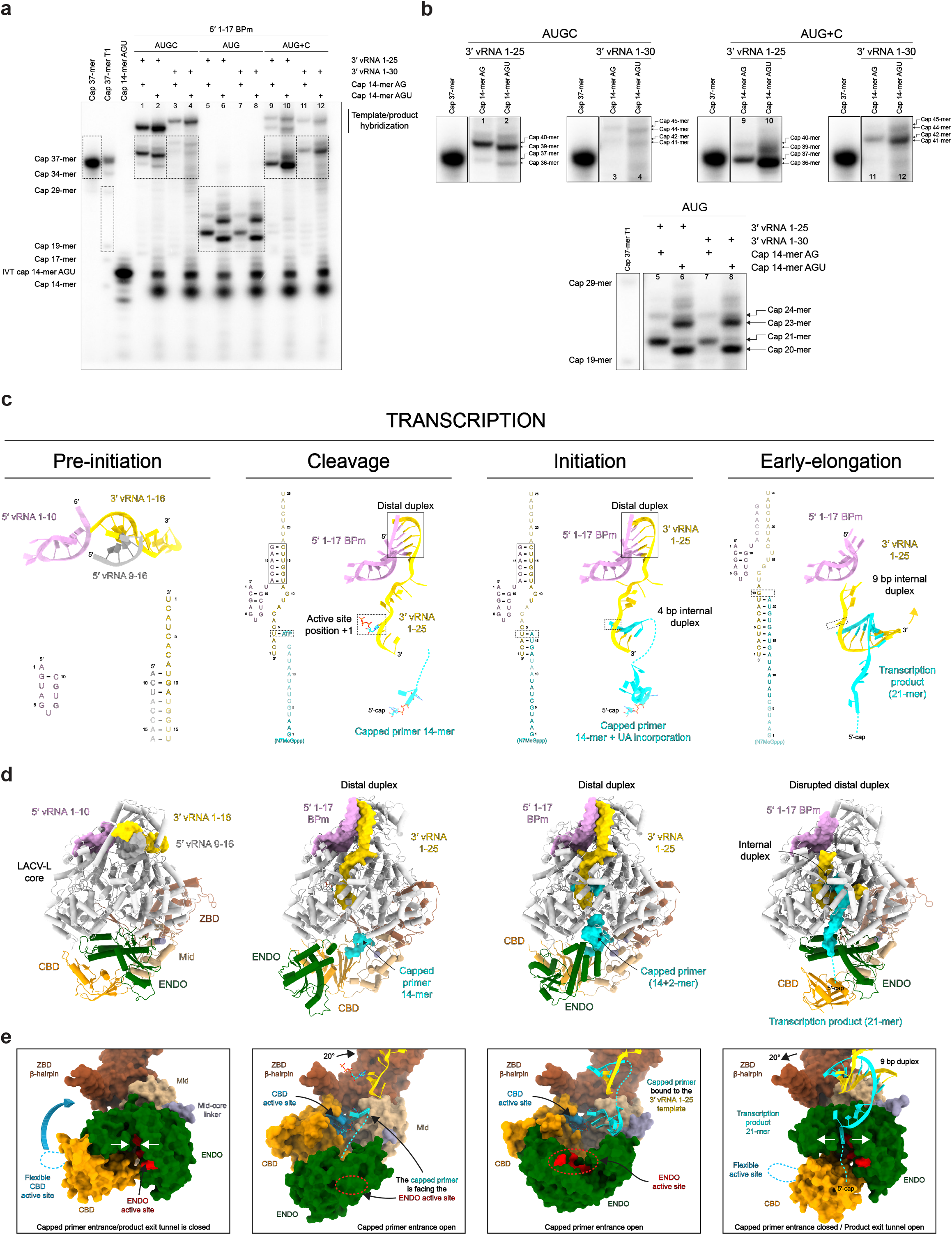
Overview of LACV-L_CItag_H34K_ activity and cryo-EM structures in transcription. **a,** *In vitro* transcription activity of LACV-L_CItag_H34K_ using different combination of 5’/3’-vRNAs, NTPs and capped primers (capped 14-mer finishing by “AG” (cap14AG) and *in vitro* produced capped 14-mer finishing by “AGU” (IVT cap14AGU)). The cap37-mer lane corresponds to a capped 37-mer identical in sequence to the theoretical transcription product of LACV-L_CItag_H34K_ with the cap14AG and the 3’-vRNA1-25 without prime-and-realign. The lane indicated as cap37-mer T1 corresponds to a RNAse T1 cleavage of the cap37. Dotted squares indicate transcription products of interest that are reported in **b**. **b,** Zoom on LACV-L_CItag_H34K_ transcription products. Lane numbers are referring to **a**. Transcription products length are indicated on the right side of the cropped gels with the corresponding molecular weight ladder. **c,** Sequence and secondary structures of 5’/3’-vRNAs, capped RNA primer and transcription products. 5’-vRNAs, 3’-vRNAs and capped primer/product are respectively colored in pink/grey, gold and cyan. Bases present in the sequences but not seen in the structures are shown in transparent. **d,** Cartoon representation of each LACV-L_CItag_H34K_ transcription structure. LACV-L domains are colored as in **Fig. 1**. RNAs are displayed as surfaces and colored as in **c**. **e,** ENDO and CBD movements during transcription. The ENDO, CBD, Mid/ZBD, RNAs are displayed as surfaces and colored as in **d**. The ENDO active site and CBD cap-binding site are colored in red and blue. The closing/opening of product exit tunnel is highlighted by white arrows.

### Overview of the transcriptionally active LACV-L structures

To visualize the path taken by the RNA during transcription initiation, a large cryo-EM dataset was collected on a transcription reaction mix containing LACV-L_CItag___H34K_, the cap14AG, the 3’-vRNA1-25, the 5’-1-17BPm, UTP, ATP and MgCl_2_. Advanced image processing enabled to separate and reconstruct several active snapshots (**Supplementary Figs. 2 and 6, Supplementary Table 2, Supplementary Movie 2**). A conformation that we suggest corresponding to the “endonuclease cleavage conformation”, captured at 3.9 Å resolution, shows the capped RNA bound to the CBD and orientated towards the ENDO cleavage site (**Fig. 4c,d**). Only the first two nucleotides of the capped RNA are visible, likely due to the low affinity of the ENDO for RNA, that is related to the diversity of sequences to be cut. The 45 Å ENDO-CBD distance is compatible with capped primer cleavage after 9 to 17 nucleotides. The 5’/3’ promoters are properly positioned for transcription initiation. The second conformation called “capped primer active site entry”, obtained at 3.1 Å resolution, shows the cap and the first two nucleotides of the capped RNA protruding towards the active site and likely corresponds to the conformation just prior transcription initiation. A “transcription initiation conformation” structure at 3.6 Å resolution is also trapped for a small-particle subset (**Fig. 4c, d**). The capped RNA is visible in the CBD and in the active site with flexible nucleotides in between. It shows incorporation of UTP and ATP in the capped RNA product. Finally, another cryo-EM data collection and image processing of the transcription complex incubated at 30° for 30 min in presence of ATP, UTP, GTP and MgCl_2_ results in the determination of a stalled early-elongation transcription conformation at 3.3 Å resolution with an elongated capped RNA that forms a 9-base pair template-product duplex in the active site cavity (**Fig. 4c, d, Supplementary Fig. 6)**.

### Major functionally relevant conformational changes of LACV-L are visualized between the transcription states

The switch from the pre-initiation to the endonuclease cleavage conformation is coupled with a large ENDO movement involving a 160° rotation that liberates the previously blocked capped RNA tunnel entrance (**Fig. 4e**). The acquired ENDO position is reminiscent to the orientation visualized in the LACV-L ΔCTER X-ray and cryo-EM structures^11^. Concomitantly to the ENDO movement, the entire CTER rotates by 20° (**Fig. 5a**), bringing the ZBD and the mid domain towards the core, where they interact with the thumb ring (**Fig. 5b**). This conformation of the CTER brings the CBD binding site close to the capped RNA entrance tunnel, contrasting with its exposed position taken at pre-initiation (**Fig. 4e**). The CBD, and in particular the loop 1850-1859 that is involved in cap binding and was disordered at pre-initiation, is stabilized by several domains and becomes visible, explaining the proper capped RNA binding site formation (**Fig. 5b**). The CBD interacts with the palm domain, the mid, the ZBD β-hairpin and the core lobe.

**Fig. 5.**
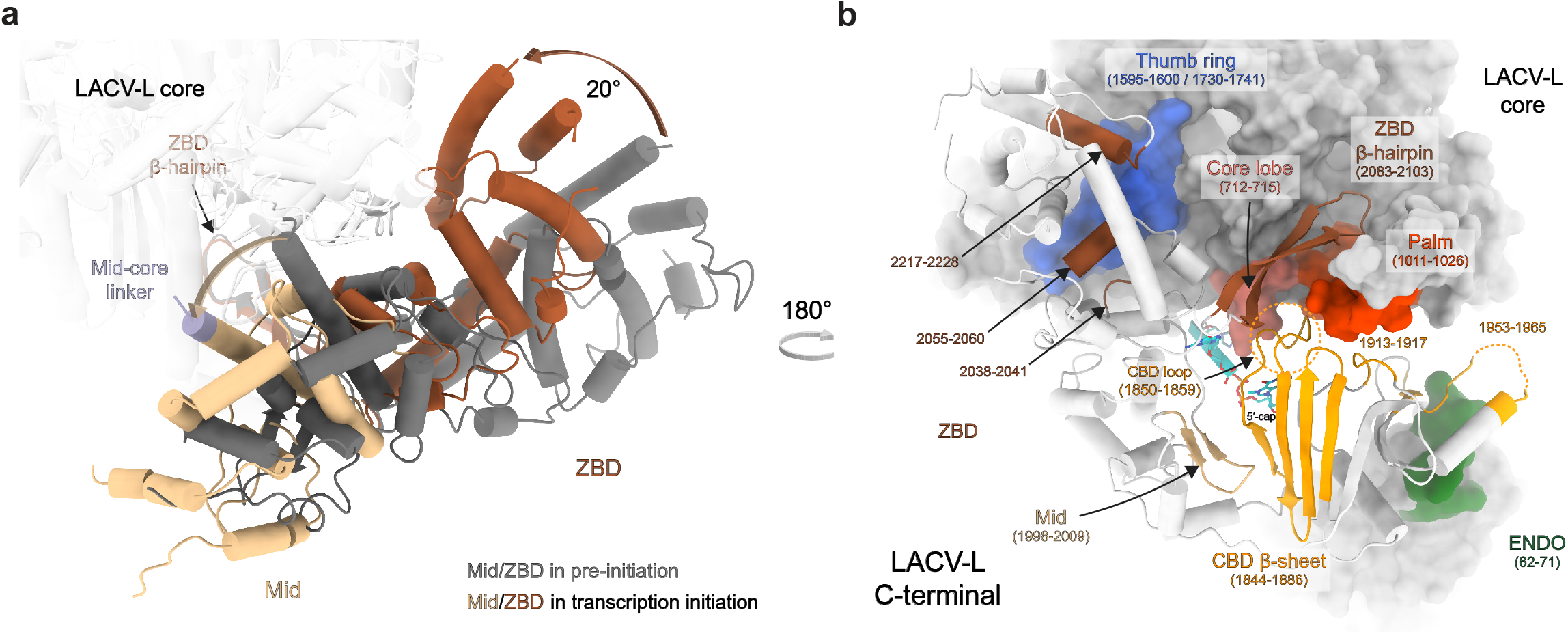
LACV-L_CItag_H34K_ conformational changes induced by transcription initiation. **a,** Mid and ZBD movements from pre-initiation to transcription initiation states. LACV-L core is transparent. Mid and ZBD are colored in grey in the pre-initiation state, in beige and brown in the transcription initiation state and their rotation highlighted with an arrow. **b,** CTER interaction with the polymerase core and the ENDO during transcription initiation. Both LACV-L core and ENDO are displayed as grey surface. The CTER domains (mid, CBD, ZBD) are displayed as cartoon. All interacting regions are displayed, colored and numbered. The CBD loop close to the CBD-binding site is surrounded by a dotted line.

Switch from the endonuclease cleavage conformation to the transcription initiation conformation involves a 175° rotation of the ENDO (**Fig. 4d, e)**. Interestingly, ENDO charge analysis suggests that, whereas the negatively charged capped RNA might be attracted by the ENDO positively charged catalytic site in the endonuclease cleavage conformation, it would be repelled by the negatively charged ENDO surface in the transcription initiation conformation, thereby facilitating capped RNA entry towards the polymerase active site (**Supplementary Fig. 7**). In addition, to the large ENDO and CTER movements, switch from pre-initiation to endonuclease cleavage conformation/transcription initiation is coupled with core closure, in an analogous manner to what is observed at replication initiation.

Nearly all the conformational changes that resulted in transcription initiation are reversed at elongation, bringing back the core in an open conformation, the ENDO and the CTER in the pre-initiation conformation (**Fig. 4d, e).** The only exception concerns the CBD that acquires an orientation differing of 30° compared to pre-initiation.

### A zoom into the key elements visualized for each transcription state reveals essential RNA-LACV-L contacts during transcription

The cleavage conformation reveals the peculiar binding of the cap that is stacked between W1847 and R1854 of the CBD, with Q1851 ensuring guanine specificity (**Fig. 6a**). The cap triphosphate moiety makes hydrogen bond interactions with K2011, K2012, R1854 and W1850. The two next nucleotides interact through their bases with Y714 of the core lobe domain, H2014, F2015, K2017 of the mid domain and Y1716 of the thumb ring domain, while the phosphate is stabilized by K2017 (**Fig. 6a**). The cleavage conformation also reveals that the PR loop tip stabilizes the template 3’ extremity prior to capped entry into the active site (**Fig. 6a**).

**Fig. 6.**
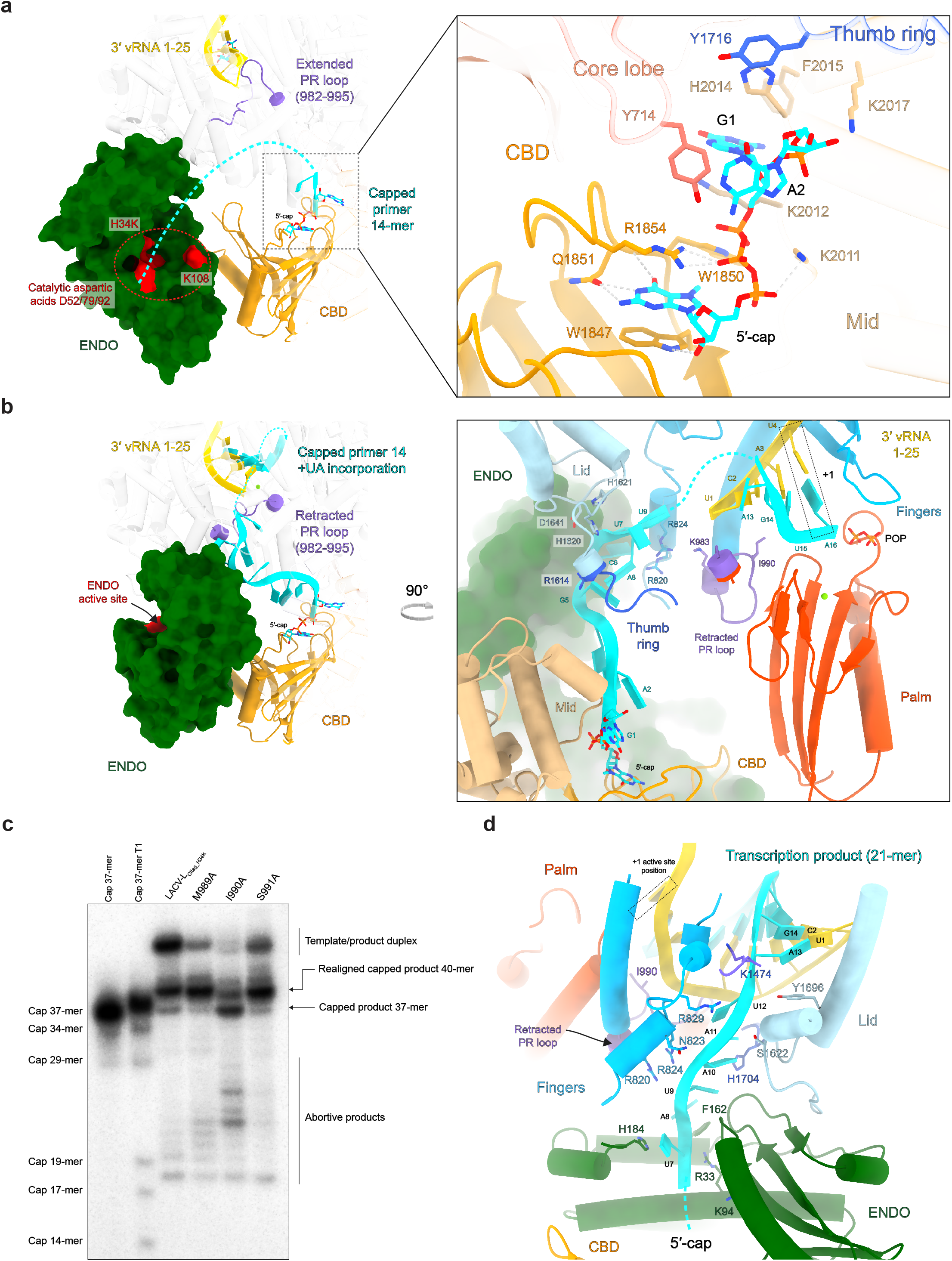
Zoom on essential RNA-LACV-L contacts during transcription. **a,** 5’-cap interaction with LACV-L_CItag_H34K_ in the transcription cleavage conformation. The ENDO active site residues are colored in red. The capped primer is colored in cyan and the dotted line stands for the potential cleavage path. On the right panel, residues implicated in the 5’-capped stabilization are displayed. **b,** Capped primer path from the CBD active site to the RdRp active site during transcription initiation. On the right panel, residues that are proximal to the capped primer are indicated. **c,** *In vitro* transcription activity of LACV-L_CItag-H34K_, LACV-L_CItag-H34K-M989A_ (M989A), LACV-L_CItag-H34K-I990A_ (I990A), LACV-L_CItag-H34K-S991A_ (S991A) with 5’-1-17BPm, cap14AG, 3’-vRNA1-25 and four NTPs. Transcription products are labelled. Cap37-mer and cap37-mer T1 lanes are equivalent to Fig. 4a description. **d,** Interaction between the capped RNA product and LACV-L_CItag_H34K_ in transcription elongation state. Residues stabilizing it are indicated. Flexible nucleotides are represented as a dotted line.

Following template binding and capped RNA cleavage, the capped RNA is directed towards the active site (**Fig. 6b**). The cap and the first 2 nucleotides of the primer remain bound in a similar fashion as in the cleavage conformation. The next nucleotides more sparsely bind through their phosphates to charged residues of the finger domain (R820, R824, K983), the thumb-ring (R1614) and the lid (H1620, H1621, D1641) (**Fig. 6b**). The binding is phosphate-specific suggesting that it is sequence independent and can probably adapt to various sizes of snatched capped RNA primer. The RNA density is more defined towards the active site, indicating a tighter binding. Capped RNA bases 13A and 14G that are complementary to 1U, 2A of the 3’-vRNA end form a duplex in position −3 and −2 of the active site. The UTP and ATP nucleotides added to the reactions were incorporated into the nascent mRNA and are positioned in position −1 and +1 of the active site in front of 3’-vRNA A3 and U4.

Following the visualized early-state initiation, a realignment must occur, as revealed by *in vitro* transcription products that are 3 nucleotides longer than if the capped RNA was simply elongated (**Fig. 4a, b**). We thus investigated the structure at initiation to identify which part of the polymerase might be implicated in realignment. Interestingly, the PR loop is in a retracted position and is proximal to the template position −4. PR loop induced template realignment from position −4 to position −1 would perfectly reposition the template for subsequent elongation (**Fig. 6b**). We thus investigated if mutation of the PR loop tip might affect realignment (**Fig. 6c**). While M989A and S991A single mutation do not affect realignment or elongation, I990A impairs realignment without preventing elongation.

Transcription then proceeds to elongation. While the template behaves like in replication, the capped RNA product exits through a tunnel that is different from the one used for its entry, surrounded by the bridge (K1474), the fingers (R820, N823, R824, R829), the lid (S1622, Y1696) and the thumb ring (H1704) (**Fig. 6d**). The capped RNA reaches the ENDO with which it interacts through residues (R33, K94, F162, H184). The 5’-cap detaches itself from the CBD, permitting the exit of capped RNA through a cleft between the ENDO and the CBD.

## DISCUSSION

### Structural elements involved in replication and transcription initiation

Proper positioning of the 3’-vRNA template in the active site is key for replication and transcription initiation. For LACV-L, it depends on (i) the distal duplex on one side and (ii) the PR loop stabilized by the ENDO on the other side. To understand the role of these elements, we describe them in light of the biochemical results acquired on Bunyaviruses and compare them with their counterparts in other sNSV polymerase structures, identifying elements conserved through all sNSV and pinpointing *Peribunyaviridae* specificities.

The observed importance of the distal duplex in proper positioning of the 3’-vRNA template is consistent with the complementarity of LACV genome that extends for 16, 21 and 23 nucleotides for the S, M and L segment respectively. Notably, complementarity of nucleotides 13 to 15 was shown to be essential for Bunyamwera, a related orthobunyavirus, in mini-replicon assays^19^. In addition, complementarity of nucleotides 16 to 19 in Bunyamwera was shown to be correlated with promoter strength, as mutation of the 3’ extremity of the S segment to make it complementary to the 5’-extremity up to nucleotide 19 resulted in an overexpressing phenotype^20^. Distal duplex formation is also observed in other sNSV and in particular Influenza virus^21^. However, 3’-vRNA template positioning for prime-and-realign, even if also used by Influenza for vRNA synthesis from cRNA^22^, has not yet been structurally visualized due to the flexibility of the 3’-cRNA extremity^23^. The LACV-L replication initiation structure thus constitutes the first of a sNSV polymerase with the distal promoter duplex compatible with prime-and-realign template positioning, posing the basis for future comparison with other sNSV polymerases that also initiate through the same mechanism.

The proper positioning of the 3’ extremity in the active site at replication also depends on the PR loop position. Although this element is not very conserved in terms of sequence within Bunyaviruses and with Orthomyxoviruses, one can see a certain conservation of the loop extremity (M989, I990, S991) in Hantavirus L proteins, which have a tyrosine, isoleucine and serine in equivalent positions (**Supplementary Fig. 8**). As M989 is likely to stabilize the incoming base in position −1, one can hypothesize that a tyrosine may play an equivalent role in hantaviruses, suggesting a conserved prime-and-realign mechanism. Interestingly, the retracted position of the PR loop is the canonical configuration of this conserved active site loop in the other structurally determined sNSV polymerases (**Supplementary Fig. 8a**). It remains to be shown whether the PR loop extension at initiation also occurs in other sNSV. Interestingly, residue V273 of influenza PB1, that corresponds to the PR loop tip, is essential for prime-and-realign both *in vitro* and *in vivo*^24^ (**Supplementary Fig. 8**). Although the influenza priming loop reaches close towards the active site and clearly plays a role in replication initiation^21,25,26^, the putative influenza PR loop may somehow complement its role.

In LACV-L, the PR loop extension at replication initiation is coupled with a conformational change of the ENDO (**Fig. 3a and Supplementary Fig. 9a**). We cannot exclude that the proper positioning of the 3’-vRNA template extremity preceding transcription initiation does not involve the same ENDO movement, given that the PR loop adopts the same position at replication initiation (**Fig. 3a**) as in the steps preceding capped RNA entry (**Fig. 6a**). In the case of influenza polymerase, although template realignment occurs following internal initiation during cRNA to vRNA replication, the situation seems to be quite different. It is thought that two distinct influenza polymerase dimers, between an apo-polymerase and the replicating polymerase, are involved in this process, whereas for LACV-L, no dimers have yet been observed. One influenza dimer has been suggested to facilitate template realignment by modifying the position of influenza priming loop^23^ (**Supplementary Fig. 9b, c**). The second replicase dimer depends essentially on the host factor ANP32, which mediates formation of dimer between a ‘packaging’ polymerase and the replicating polymerase^27^. Both influenza polymerases in both these dimers are in the ‘replicase’ rather than ‘transcriptase’ conformation, which involves a re-positioning of the ENDO and re-organisation of the PB2-CTER that brings the PB2-cap and the residues 490-493 of the Cap-627 linker close to the putative PR loop (**Supplementary Fig. 9d)**. It remains to be seen if the residues 490-493 of the Cap-627 linker and the CBD may, when acquiring this position, trigger a rearrangement of the PR loop that could complement the role of the priming loop in the prime-and-realign mechanism.

Proper transcription initiation, on top of the 3’ template proper positioning, depends on capped RNA binding, cleavage and positioning in the active site. In LACV-L, major conformational changes of the ENDO and the CTER are involved in this process. These movements are more pronounced than for influenza polymerase, in which the ENDO remains in the same position between the cleavage and the transcription initiation conformations, the only major movement being done by the PB2-cap that rotates of 70° to bring the capped RNA first to the ENDO active site and then to the polymerase active site^21,28^. The CBD and ENDO of LACV and influenza polymerases are in different positions compared to the core precluding their exact comparison but the distance separating the cap and the ENDO cleavage site are of 45 Å for LACV and 49/52 Å for Influenza, in accordance with the similar sizes of capped primers (**Supplementary Fig. 10**). Although the CBD positions differ, the capped RNA tunnel entrance is conserved, together with core closing upon initiation. However, the details of transcription initiation differ in the two polymerase due to the exact positioning of the 3’-template in the active site at initiation, with the first nucleotide of the vRNA template being in position −2 and −3 for Influenza and LACV respectively^25^ (**Supplementary Fig. 10**).

### Proposed models for replication and transcription initiation by prime-and-realign

Combination of *in vitro* activity assays and structural snapshots of LACV-L in different active conformations lead us to propose a structural model for the prime-and-realign mechanism. Concerning replication, we hypothesize that proper closing of the active site motifs coupled with the visualized flexibility of the nucleotides 6, 7, 8 of the 3’ template enable a transient translocation of both the template and ATP of 1 nucleotide, thereby positioning the ATP in position −1 of the active site, where it would interact with the residue M989 of the PR loop tip (**Fig. 7b**). This would enable positioning of a GTP in position 1 of the active site and formation of a nascent pppApG product. Normal elongation would generate a pppApGpU product concomitantly with translocation of the template with its extremity located in position −5 of the active site. At this stage, the template would be in extreme tension due to (i) the presence of the distal duplex linked to the 5’-hook at the tunnel entrance, (ii) the maximum stretching of template nucleotides 6-8 and (iii) the template positioning constrains by the PR loop. These would act as a spring and break the hydrogen bonds between pppApGpU and the template nucleotide 4 to 6, repositioning the primer in front of template nucleotide 1 to 3, completing the realignment. Further elongation results in proper product formation, exactly complementary to the template.

**Fig. 7.**
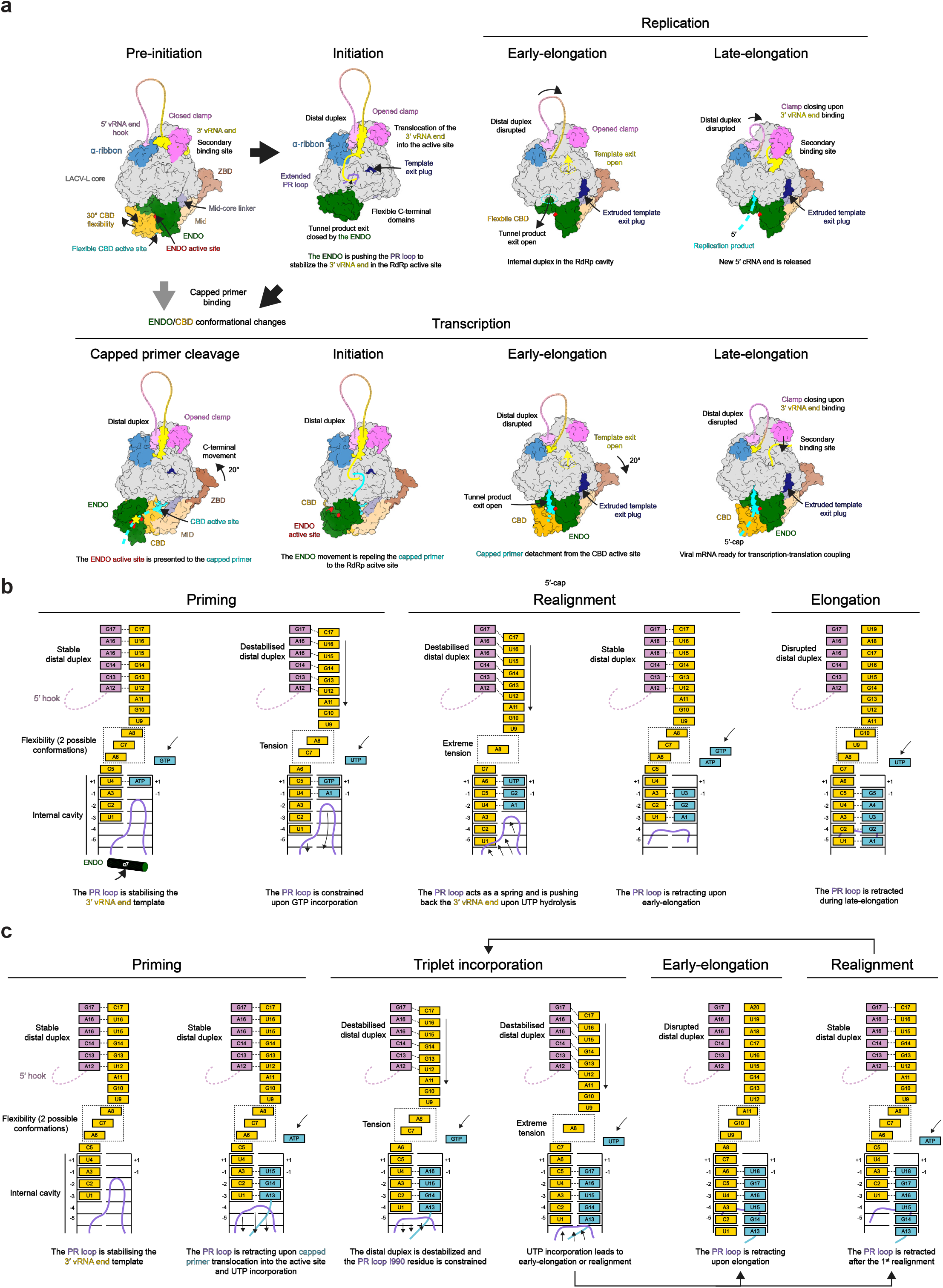
LACV-L replication and transcription cycle models. **a,** Structure-based model of the mechanisms underlying LACV-L replication and transcription. All structures are displayed as surfaces with specific domains and features colored as in **Fig. 1 and 2**. The vRNA is shown as a line with the 5’/3’ extremities colored in pink/yellow. At initiation, the 3’-vRNA extremity is released from the secondary binding site and brought into the RdRp active site where it is stabilized by the PR loop, that extends due to the ENDO movement. If the flexible CBD snatches a 5’-cap, important conformational changes of the CBD, the ENDO, the mid and the ZBD occur, triggering capped primer translocation into the ENDO active site (in red). Once cleaved (yellow star), an ENDO movement triggers the capped primer repositioning towards the RdRp active site. Early elongation induces extrusion of the template exit plug resulting in the template exit opening (yellow dotted circle). In late-elongation, the 3’-vRNA template extremity goes back to the secondary binding site. **b,c,** Structure-based model of initiation by prime-and-realign for replication **(b)** and transcription **(c)**. The 5’/3’-vRNA ends are respectively colored in pink and gold. The 5’-hook structure is represented as a dotted line. Incorporated nucleotides are colored in blue. The PR loop is shown in pink. The binding pocket of nucleotides A6, C7 and A8 at priming stage is surrounded by a dotted square. The proposed successive steps of the models are presented from left to right. The main role of each step is indicated below the each schematic.

Concerning transcription, it is expected that the template and capped RNA positioning are more permissive than in replication, to adapt to the capped RNA length and composition (**Fig. 7c**). Capped RNA terminating by AG would position in front of the 3’ template extremity in position −2 and −1 of the active site, ready for incorporation of a U in position 1. This would require the retraction by one nucleotide of the template, made possible by the flexibility of its nucleotides 6-8 and capped RNA hybridization. This would permit incorporation of UTP in position +1 of the active site. Subsequent translocation would bring the polymerase in the same state as if a capped RNA terminating by AGU is used as a primer. The 3’ template extremity would be in position −3 to −1 of the active site, ready for incorporation of an A in position 1. Following incorporation of the subsequent UTP and GTP, realignment may occur once or several times, accordingly to *in vivo* mRNA composition observation. This may be facilitated by the PR loop, which, by interfering with proper hydrogen bonding between the template extremity and the product, may reduce the number of hydrogen bonds to be broken upon realignment. Although speculative, this is supported by the absence of interaction of the extreme 3’ A nucleotide of the template with its complementary U of the capped RNA in position −3 of the active site (**Fig. 6b**). As for replication, we propose that (i) the destabilization of the distal duplex, (ii) the extreme tension of nucleotides 6-8, (iii) and the PR constrains may provide enough energy to trigger template repositioning. The residue I990 of the PR loop is likely to be involved in pushing back the template from position −4 to position −1 of the active site as evidenced by transcription assay done in presence of I990A mutant (**Fig. 6c**). The dynamic of the process, that involves a large conformational change from initiation to elongation (**Fig. 4c-e**) is key in the realignment, as evidenced by the diminution of realignment for reactions initially done with three nucleotides and subsequently supplemented with CTP (**Fig. 4a, b lines 9-12**). Following realignment(s) (if realignment occurs) the PR loop would retract, thereby switching the polymerase to an elongation mode.

The proposed models for replication and transcription prime-and-realign may provide hints on how these processes may occur in other families of the *Bunyavirales* order^4–6,29,30^ and complements the existing knowledge on prime-and-realign mechanisms used by Influenza for the replication of vRNA from a cRNA template^22,24^, and for transcription^31^.

In conclusion, the combination of *in vitro* assays and cryo-EM structural snapshots of specific LACV-L active states unveils the precise mechanisms of initiation by prime-and-realign and elongation of both replication and transcription. These results are novel for the entire *Bunyavirales* order and shed light on processes conserved in all sNSV. They will be key to design structure-based drugs to counteract these life-threatening viruses that would target essential elements of replication and transcription activity or prevent essential conformational changes.

## METHODS

### LACV minigenome assay

Sub-confluent monolayers of HEK-293 cells seeded in 6-well dishes were transfected with 250 ng each of plasmid pHH21-LACV-vMRen, pTM-LACV_L (or insertion mutants), pTM-LACV_N, 500 ng pCAGGS-T7, and 100 ng pTM1-FF-Luc (described in^32,33^ using Nanofectin transfection reagent in 200 μl serum-free medium (OptiMEM, Gibco-BRL). In addition, 250 ng of pI.18-HA-PKR was added to the plasmid mix^32^. At 72 h post-transfection, cells were lysed in 200 ml Dual Luciferase Passive Lysis Buffer (Kit Promega) and both Luciferase (FF-Luc) and Renilla luciferase (REN-Luc) activities were measured as described by the manufacturer (Promega). As pTM-LACV_L, we used N-terminal (Nter) His-tag LACV-L full-length construct (strain LACV/mosquito/1978, GenBank: EF485038.1, UniProt: A5HC98) previously cloned into a pFastBac1 vector between NdeI and NotI restriction sites (Arragain et al, 2020), Cterminal (Cter) His-tag LACV-L full length construct. An insertion mutant of LACV-L full length construct (insertions of 2 internal Strep-Tag II “WSHPQFEK” flanked by a SG/GSG linkers) was as well generated between residues G1034-Y1035.

### Cloning, expression and purification

The LACV-L full length construct with an addition of one internal Strep-Tag II “WSHPQFEK” flanked by a SG/GSG linker between residues G1034 and Y1035 (G1034-SG-StrepTag II-GSG-Y1035) was chosen and recloned. An additional H34K mutation was inserted leading to the LACV-L_CItag_H34K_ construct used for cryo-EM analysis and activity assays. Mutations M989A/I990A/S991A (LACV-L_CItag_H34K_M989A_/LACV-L_CItag_H34K_I990A_/LACV-L_CItag_H34K_S991A_) were inserted on LACV-L_CItag_H34K_ construct and used for activity assays testing the role of the PR loop. All LACV-L clones were made using combination of Polymerase Chain Reactions (PCR), agarose gel purification, DNA extraction using PCR clean-up kit (Machery-Nagel), Gibson assembly (NEB) and finally sequenced (Genewiz) before proceeding to further experiments. For all LACV-L_CItag_H34K_ constructs, expressing baculoviruses were made using the Bac-to-Bac method (Invitrogen)^34^. For protein expression, High 5 (Hi5) cells at 0.5 × 10^6^ cells/mL concentration were infected by adding 0.1% of virus and collected 72 h to 96 h after the day of proliferation arrest.

Hi5 cells were resuspended in lysis buffer (50 mM Tris–HCl pH 8, 500 mM NaCl, 2 mM β-mercaptoethanol (BME), 5% glycerol) with cOmplete EDTA-free protease inhibitor cocktail (Roche) and disrupted by sonication for 3 min (10 s ON, 20 s OFF, 50% amplitude) on ice. Lysate was clarified by centrifugation at 48,000 g during 45 min at 4 °C and filtered. Soluble fraction was loaded on pre-equilibrated StrepTrap HP column (Sigma-Aldrich) and eluted using initial lysis buffer supplemented by 2 mM d-Desthiobiotin (Sigma-Aldrich). LACV-L_CItag_H34K_ fractions were subsequently pooled, dialyzed 1h at 4 °C in heparin buffer (50 mM Tris–HCl pH 8, 250 mM NaCl, 2 mM BME, 5% glycerol) and loaded on HiTrap Heparin HP column (Sigma-Aldrich). Elution was performed using 50 mM Tris–HCl pH 8, 1 M NaCl, 5 mM BME, 5% glycerol. LACV-L_CItag_H34K_ protein was finally loaded on a pre-equilibrated Superdex 200 Increase 10/300 GL (Sigma-Aldrich) size exclusion chromatography column in 50 mM Tris–HCl pH 8, 400 mM NaCl, 5 mM BME. Best fractions were pulled, concentrated using Amicon Ultra 10 kDa (Merck Millipore), flash frozen in liquid nitrogen and conserved at −80°C for future experiments.

### *In vitro* transcription, capping and RNAse T1 cleavage

*In vitro* transcription was realized using a DNA oligo enclosing the T7 promoter sequence with an additional G at the 3’ end (5’-TAA TAC GAC TCA CTA TAG <u>G</u>-3’) and a template DNA oligo enclosing both complementary (i) T7 promoter sequence and (ii) 3’-5’ DNA sequence of the desired 5’-3’ RNA.

Two different templates were used to produce a 37-mer RNA (3’-ATT ATG CTG AGT GAT ATC <u>C</u>TA CGA TAT TAT CAT CAC ATG ATG GTT CAT ATC TAT-5’ giving 5’-G<u>G</u>A UGC UAU AAU AGU AGU GUA CUA CCA AGU AUA GAU A-3’) and a 14-mer RNA (3’-ATT ATG CTG AGT GAT ATC <u>C</u>TA CGA TAT ATC A-5’ giving 5’-G<u>G</u>A UGC UAU AUA GU-3’).

T7 promoter (15 μM) and template DNA (10 μM) oligos were mixed in transcription buffer (30 mM TRIS-HCl pH 8, 20 mM MgCl_2_, 0.01% Triton X-100, 2 mM Spermidine), annealed at 80°C for 2 min and cooled down to room temperature (RT). Hybridized DNA oligos (1 μM) were then added to transcription buffer supplemented by 5 mM ATP, UTP, GTP and CTP (Sigma-Aldrich), 10 mM dithiothreitol (DTT), 1% PEG8000, 50 μg/mL T7 RNA polymerase and incubated overnight at 37°C. *In vitro* transcription reactions were stopped by adding 2X RNA loading dye (95% formamide, 1 mM EDTA, 0.025% SDS, 0.025% bromophenol blue, 0.01% xylene cyanol), heated 5 min at 95°C and loaded on a 20% Tris-Borate-EDTA-7M urea-polyacrylamide gel. The corresponding products bands were cut, soaked in 0.3 M NaOAc pH 5.0-5.2 and put at rocking overnight at 4°C. After filtration, RNAs solutions were supplemented with 3V of 100% ethanol, vortexed and put 2h at −80°C. Precipitated RNAs were pelleted by centrifugation at 48,000 g during 20 min at 4°C. The supernatant was discarded and RNAs pellets were carefully washed with 70% ethanol before another centrifugation at 48,000 g during 20 min at 4°C. After two consecutive washes, 70% ethanol was removed, RNAs pellets were dried 10 to 15 min before being resuspended in 50 mM TRIS-HCl pH 8 and stored at −20°C.

*In vitro* RNA capping was achieved using RNAs produced by *in vitro* transcription as substrate and the Vaccinia capping system (NEB). The 37-mer (50 μM) and the 14-mer (20 μM) RNAs were mixed with Vaccinia Capping Enzyme buffer (NEB), 2 mM SAM, 3 pM GTP α-^32^P (PerkinElmer), 10 Units of VCE and incubated 30 min at 37°C.

To produce capped molecular markers, 2X RNA loading dye was added to capped RNA and solution was heated 5 min at 95°C before loading on a 20% TBE-Urea-Polyacrylamide gel. Visible band corresponds to capped 37-mer (5’-m7Gppp G<u>G</u>A UGC UAU AAU AGU AGU GUA CUA CCA AGU AUA GAU A-3’) and capped 14-mer (5’-m7Gppp G<u>G</u>A UGC UAU AUA GU-3’). Additionally, capped 37-mer was digested using RNase T1 cleavage kit (Ambion). Cleavage reaction was performed for 15 min at 37°C, stopped by adding 2X RNA loading dye and heating 5 min at 95°C. Observed bands corresponds to a 34-mer, a 29-mer, a 19-mer, a 17-mer and a 14-mer RNA (5’-m7Gppp GGA UGC UAU AAU AG/U AG/U G/UA CUA CCA AG/U AUA G/AU A-3’).

### Replication and transcription activity assay

Synthetic RNAs (Microsynth) 5’-vRNA1-17 (5’-AGU AGU GUG CUA CCA AG-3’), 5’-1-17BPm (5’-A<u>CG</u> AGU GU<u>C</u> <u>G</u>UA CCA AG-3’), 3’-vRNA1-25 (5’-UAU CUA UAC UUG GUA GUA CAC UAC U-3’), 3’-vRNA1-30 (5’-AAC GUU AUC UAU ACU UGG UAG UAC ACU AC U-3’) were used as viral RNAs (vRNA). Capped 14-mer ending by “AG” (5’-m7Gppp GAA UGC UAU AAU AG-3’) (TriLink Biotechnologies) and *in vitro* produced capped 14-mer ending by “AGU” (5’-m7Gppp GGA UGC UAU AUA GU-3’) were used as capped primers.

For replication activity assays, 0.6 μM LACV-L_CItag_H34K_ were mixed with (i) 0.9 μM 5’-vRNA for 1h at 4°C, then 0.9 μM 3’-vRNA for overnight incubation at 4°C.

Reactions were launched at RT, 27°C, 30°C, 37°C for 0.5, 1, 2, 3 and 4h by adding all NTPs or only ATP/GTP/UTP at 100 μM/NTP, 0.75 μCi/ml α-^32^P GTP and MgCl_2_/MnCl_2_ in a final assay buffer containing 50 mM TRIS-HCl pH 8, 150 mM NaCl, 5 mM BME, 100 μM/NTP, 0-1-2-3-4-5 mM MgCl_2_/MnCl_2_. Decade markers system (Ambion) was used as molecular weight ladder. For transcription activity assays, same conditions were used as in replication activity assay except that 0.9 μM capped RNA primer was added at the same time as the 5’-1-17BPm. The 37-mer capped primer, before and after RNAse T1 cleavage, was used as molecular weight ladder.

Reactions were stopped by adding 2X RNA loading dye, heating 5 min at 95°C and immediately loaded on a 20% TBE-7M urea-polyacrylamide gel that was run 2h at 20 W. The gel was exposed on a storage phosphor screen and read with an Amersham Typhoon scanner.

### Electron microscopy

To catch structural snapshots of LACV-L_CItag_H34K_ in replication early-elongation state, 1.3 μM LACV-L_CItag_H34K_ were sequentially incubated for 1h at 4°C with (i) 2.6 μM 5’-1-17BPm, (ii) 2.6 μM 3’-vRNA1-25. LACV-L_CItag_H34K_ bound to vRNAs was subsequently incubated for 4h at 30°C in a final buffer containing 50 mM TRIS-HCl pH 8, 150 mM NaCl, 5 mM MgCl_2_, 5 mM BME and 100 μM of each NTP (ATP/GTP/UTP).

To catch structural snapshots of LACV-L_CItag_H34K_ in replication late-elongation, 1.7 μM LACV-L_CItag_H34K_ were sequentially incubated for 1h at 4°C with (i) 1.9 μM 5’-1-17BPm, (ii) 1.9 μM 3’-vRNA1-30. LACV-L_CItag_H34K_ bound to vRNAs was subsequently incubated for 4h at 30°C in a final buffer containing 50 mM TRIS-HCl pH 8, 150 mM NaCl, 5 mM MgCl_2_, 5 mM BME and 100 μM of each NTP (ATP/GTP/UTP/CTP).

To catch structural snapshots of LACV-L_CItag_H34K_ in transcription, 1.3 μM LACV-L_CItag_H34K_ were sequentially incubated for 1h at 4°C with (i) 1.9 μM 5’-1-17BPm and 3.9 μM commercial 14-mer capped primer finishing by “AG”, (ii) 1.9 μM 3’-vRNA1-25. LACV-L_CItag_H34K_ bound to vRNAs and capped primer was subsequently incubated for 1h at 30°C in a final buffer containing 50 mM TRIS-HCl pH 8, 150 mM NaCl, 2 mM MgCl_2_, 5 mM BME and 100 μM of NTPs (ATP/UTP) for transcription cleavage/capped primer entry/initiation states or 100 μM of each NTP (ATP/GTP/UTP) for transcription early-elongation state.

Both replication and transcription reaction were put at 4°C before grids freezing. For each reaction, 3.5 μl of sample were deposited on negatively glow-discharged (25 mA, 45 sec) UltraAuFoil gold grids 300 mesh, R 1.2/1.3. Excess solution was blotted for 2 sec, blot force 1, 100% humidity at 20°C with a Vitrobot Mark IV (Thermo Fischer Scientific) before plunge-freezing in liquid ethane.

Automated data collections of (i) the replication early-elongation state, (ii) the replication late-elongation state, (iii) the transcription initiation state and (iv) the transcription early-elongation state were performed on a 200 kV Glacios cryo-TEM microscope (Thermo Fischer Scientific) equipped with a K2 direct electron detector (Gatan) using SerialEM^35^. Coma and astigmatism correction were performed on a carbon quantifoil grid. Movies of 60 frames were recorded in counting mode at a 36,000× magnification giving a pixel size of 1.145 Å with defocus ranging from −0.8 to −2.0 μm. Total exposure dose was 60 e^−^/Å^2^. The number of movies per experiment are indicated in **Supplementary Figs. 2, 3 and 6** and in **Supplementary Tables 1 and 2**.

To improve the resolution of the (i) replication initiation state, the (ii) transcription capped primer entry state and the (iii) transcription initiation state, an automated data collection was performed on a Titan Krios G3 (Thermo Fischer Scientific) operated at 300 kV equipped with a K3 (Gatan) direct electron detector camera and a BioQuantum energy filter using SerialEM. Micrographs were recorded in counting mode at a 130,000× magnification giving a pixel size of 0.645 Å with defocus ranging from −0.8 to −1.8 μm. Movies of 40 frames were collected with a total exposure of 50 e^−^/Å^2^. 15573 movies were collected.

### Image processing

For each collected dataset, movie drift correction was performed in Motioncor2^36^. For images collected on the Thermo Fischer Scientific Glacios TEM, the two first frames and the last ten were removed. For images collected on the Thermo Fischer Scientific Titan Krios TEM, only the first two first frames were removed. Both gain reference and camera defect corrections were applied. Further initial image processing steps were performed in cryoSPARC v3.2.0^37^. CTF parameters were determined using “Patch CTF estimation” on non-dose weighted micrographs. Realigned micrographs were then inspected and low-quality micrographs displaying crystalline ice, ice contamination or aggregates for example were manually discarded for further image processing. LACV-L_CItag_H34K_ particles were picked using a circular blob with a diameter ranging from 90 to 170 Å. The number of particles picked is indicated in **Supplementary Figs. 2, 3 and 6** and in **Supplementary Tables 1 and 2**. For Thermo Fischer Scientific Glacios datasets, particles were extracted from dose-weighted micrographs using a box size of 260 × 260 pixels^2^. For the Thermo Fischer Scientific Titan Krios dataset, the box size was 440 × 440 pixels^2^. For each dataset, the same image processing approach was used to separate the different LACV-L conformation and get the best map quality. First, successive 2D classifications were used to eliminate bad quality particles displaying poor structural features. LACV-L_CItag_H34K_ initial 3D reconstruction was generated with “Ab-initio reconstruction” in cryoSPARC using a small subset of particles. All selected particles were subjected to 3D refinement with per-particle CTF estimation. The rest of the image processing was done in RELION 3.1^38,39^. For each dataset, particles were divided in equal subset of 250k particles and subjected to multiple 3D classification (10 classes for each subset) with coarse image-alignment sampling using a circular mask of 170 Å. For each identical LACV-L conformation, particles were grouped and subjected to a 3D refinement with a circular mask of 170 Å followed by 3D classification using local angular searches based on previously determined orientations. High-resolution classes and particles belonging to the same conformation were finally grouped and subjected to 3D masked refinement with local angular searches to obtain high-resolution structure. Masks to perform sharpening were generated using last 3D refined maps which has been low-pass filtered at 10 Å, extended by 4 pixels with 8 pixels of soft-edge. Post-processing was done using a manually determined *B*-factor. For each final map, reported global resolution is based on the FSC 0.143 cut-off criteria and local resolution variations were also estimated in RELION 3.1 (**Supplementary Figs. 2, 3 and 6**).

### Model building in the cryo-EM maps

Both LACV-L in pre-initiation state (PDB: 6Z6G) and LACV-L in elongation mimicking state (PDB: 6Z8K) models^12^ were used as a starting point to manually build into the different cryo-EM maps in either replication/transcription initiation/elongation states using COOT^40^. All the RNAs (5’-1-17BPm, 3’-vRNA1-25, 3’-vRNA1-30, the capped primer and both replication and transcription products) were manually built using COOT. Each model was refined using Phenix-real space refinement^41^. Atomic model validation was performed using the Phenix validation tool and the PDB validation server. Model resolution according to cryo-EM map was estimated at the 0.5 FSC cutoff. Figures were generated using ChimeraX^42^. Electrostatic potential was calculated using PDB2PQR^43^ and APBS^44^.

### Multiple alignment

Multiple alignment was performed using Muscle^45^ and is displayed using ESPript^46^.

## DATA AVAILABILITY

Coordinates and structure factor have been deposited in the Protein Data Bank and the Electron Microscopy Data Bank.

LACV-L replication initiation state PDB 7ORN EMDB EMD-13043

LACV-L replication early-elongation state PDB 7ORO EMDB EMD-13044

LACV-L replication late-elongation state PDB 7ORI EMDB EMD-13038

LACV-L transcription capped primer cleavage state PDB 7ORJ EMDB EMD-13039

LACV-L transcription capped primer active site entry state PDB 7ORK EMDB EMD-13040

LACV-L transcription initiation state PDB 7ORL EMDB EMD-13041

LACV-L transcription early-elongation state PDB 7ORM EMDB EMD-13042

## ACKNOWLEDGEMENTS

We thank Karine Huard, Angélique Fraudeau, Petra Drncová, Alice Aubert and Martin Pelosse for technical advices on expression and purification; Friedemann Weber for providing the LACV mini-replicon to S.C.; Sissy Kalayil, Tomas Kouba, Anna Dubankova and Joanna Wandzik for technical advices on radioactivity experiments; Lionel Imbert and Alice Stelfox for advices on *in vitro* transcription; Simon Fromm for data collection on the Heidelberg Titan Krios; Aymeric Peuch for setting up and maintaining the EM computing cluster; Piotr Gerlach and Juan Reguera for useful discussions and Daphna Fenel for technical support.

This work used the platforms of the Grenoble Instruct-ERIC center (ISBG; UAR 3518 CNRS-CEA-UGA-EMBL) within the Grenoble Partnership for Structural Biology (PSB), supported by FRISBI (ANR-10-INBS-05-02) and GRAL, financed within the University Grenoble Alpes graduate school (Ecoles Universitaires de Recherche) CBH-EUR-GS (ANR-17-EURE-0003). The electron microscope facility is supported by the Auvergne-Rhône-Alpes Region, the Fondation pour la Recherche Médicale (FRM), the fonds FEDER and the GIS-Infrastructures en Biologie Santé et Agronomie (IBISA). This work benefited from access to the cryo-electron microscopy platform of the European Molecular Biology Laboratory (EMBL) in Heidelberg and has been supported by iNEXT-Discovery, project number 871037, funded by the Horizon 2020 program of the European Commission. We thank all platform staff that enabled us to perform these analyses. IBS acknowledges integration into the Interdisciplinary Research Institute of Grenoble (IRIG, CEA).

This work was supported by the ANR-19-CE11-0024-02 and the Institut Universitaire de France endowment to H.M., the FRM end of PhD grant to B.A.

## AUTHOR CONTRIBUTIONS

F.B. and B.A. cloned LACV-L FL with internal tags. F.B. performed mini-replicon assays. B.A. and Q.D.T. cloned mutant constructs. B.A. and Q.D.T. expressed and purified LACV-L. B.A. performed *in vitro* activity assays. B.A. and H.M. prepared cryo-EM grids. H.M. and B.A. collected cryo-EM data on a Thermo Fischer Scientific Glacios EM thanks to advices and training from G.S. who set up and maintains the IBS-ISBG EM platform. B.A. performed cryo-EM image processing. B.A and H.M. built the models based on the cryo-EM maps with inputs from S.C. B.A. and H.M. performed structural analysis. H.M. and G.S. co-supervise B.A. and Q.D.T. The project was conceived by H.M. with inputs from S.C. and G.S. This project used funding obtained by H.M. and G.S. The manuscript was written by H.M. and B.A. with input from all authors.

