## Supplementary Tables 1 and 2, Supplementary Figures 1 to 10 for "Structural snapshots of La Crosse virus polymerase reveal the mechanisms underlying *Peribunyaviridae* replication and transcription"

**Benoît Arragain *et al.***

**Supplementary Table 1. Cryo-EM data collection, refinement and validation statistics of LACV-L replication snapshots**

| | **LACV L Replication initiation**<br><br>(PDB 7ORN)<br>(EMD-13043) | **LACV L Replication early elongation**<br><br>(PDB 7ORO)<br>(EMD-13044) | **LACV L Replication late-elongation**<br><br>(PDB 7ORI)<br>(EMD-13038) |
|---|---|---|---|
| **Data collection and processing** | | | |
| Microscope | Thermo Fisher Scientific Titan Krios | Thermo Fisher Scientific Glacios | Thermo Fisher Scientific Glacios |
| Camera | Gatan K3 | Gatan K2 Summit | Gatan K2 Summit |
| Magnification | 130000 | 36000 | 36000 |
| Voltage (kV) | 300 | 200 | 200 |
| Number of frames | 40 | 60 | 60 |
| Electron exposure ($e^-/A^2$) | 50 | 60 | 60 |
| Defocus range (μm) | -0.8 to -1.8 | -0.8 to -2.0 | -0.8 to -2.0 |
| Pixel size (Å) | 0.645 | 1.145 | 1.145 |
| Symmetry imposed | C1 | C1 | C1 |
| Initial/Final micrographs (no.) | 15573/15444 | 2927/2923 | 1848/1841 |
| Final particles (no.) | 641008 | 417757 | 24151 |
| Map resolution (Å) 0.143 FSC threshold | 2.8 | 2.9 | 3.9 |
| Map resolution range (Å) | 2.6-3.5 | 2.7-4 | 3.6-6 |
| **Refinement** | | | |
| Initial model used | 6Z6G | 6Z8K | 6Z8K |
| Model resolution (Å) 0.5 FSC threshold | 2.8 | 3.0 | 3.9 |
| Map sharpening B factor ($Å^2$) | -50 | -50 | -50 |
| Model composition | | | |
| Protein residues | 1700 | 2008 | 2007 |
| Nucleotide residues | 35 | 48 | 44 |
| Ligands | 3 | 2 | 3 |
| Water | 78 | 0 | 0 |
| B-factor ($Å^2$, min-max (mean)) | | | |
| Protein | 38.17-142.20 (64.15) | 37.78-160.62 (74.97) | 68.43-192.36 (109.75) |
| Nucleotides | 44.49-108.37 (67.92) | 51.26-142.63 (75.16) | 97.23-188.85 (112.03) |
| Ligands | 51.03-53.69 (51.85) | 46.51-133.86 (90.19) | 88.19-172.55 (112.03) |
| R.m.s deviations | | | |
| Bond lengths (Å) | 0.003 | 0.004 | 0.003 |
| Bond angles (°) | 0.508 | 0.531 | 0.501 |
| Validation | | | |
| MolProbity score | 1.37 | 1.52 | 1.69 |
| Clashscore | 5.27 | 6.40 | 9.32 |
| Poor rotamers (%) | 0.06 | 0.05 | 0.00 |
| Ramachandran plot | | | |
| Favored (%) | 97.57 | 97.04 | 96.79 |
| Allowed (%) | 2.43 | 2.96 | 3.21 |
| Disallowed (%) | 0 | 0 | 0 |

**Supplementary Table 2. Cryo-EM data collection, refinement and validation statistics of LACV-L transcription snapshots**

| | **LACV L Transcription cleavage conformation** (PDB 7ORJ) (EMD-13039) | **LACV L Capped primer active site entry** (PDB 7ORK) (EMD-13040) | **LACV L Transcription initiation** (PDB 7ORL) (EMD-13041) | **LACV L Transcription early elongation** (PDB 7ORM) (EMD-13042) |
|---|---|---|---|---|
| **Data collection and processing** | | | | |
| Microscope | Thermo Fisher Scientific Glacios | Thermo Fisher Scientific Titan Krios | Thermo Fisher Scientific Titan Krios | Thermo Fisher Scientific Glacios |
| Camera | Gatan K2 Summit | Gatan K3 | Gatan K3 | Gatan K2 Summit |
| Magnification | 36000 | 130000 | 130000 | 36000 |
| Voltage (kV) | 200 | 300 | 300 | 200 |
| Number of frames | 60 | 40 | 40 | 60 |
| Electron exposure ($e^-/A^2$) | 60 | 50 | 50 | 60 |
| Defocus range (μm) | -0.8 to -2.0 | -0.8 to -1.8 | -0.8 to -1.8 | -0.8 to -2.0 |
| Pixel size (Å) | 1.145 | 0.645 | 0.645 | 1.145 |
| Symmetry imposed | C1 | C1 | C1 | C1 |
| Initial/Final micrographs (no.) | 3270/3149 | 15573/15444 | 15573/15444 | 2524/2363 |
| Final particles (no.) | 29065 | 76579 | 19112 | 229480 |
| Map resolution (Å) 0.143 FSC threshold | 3.9 | 3.1 | 3.6 | 3.3 |
| Map resolution range (Å) | 3.6-6.0 | 2.8-5 | 3.3-5 | 3.1-5.5 |
| **Refinement** | | | | |
| Initial model used | 6Z6G | 6Z6G | 6Z6G | 6Z8K |
| Model resolution (Å) 0.5 FSC threshold | 3.9 | 3.0 | 3.6 | 3.4 |
| Map sharpening B factor ($Å^2$) | -60 | -50 | -50 | -80 |
| Model composition | | | | |
| Protein residues | 2129 | 2182 | 2183 | 2145 |
| Nucleotide residues | 36 | 37 | 47 | 40 |
| Ligands | 2 | 4 | 3 | 2 |
| Water | 0 | 48 | 0 | 0 |
| B-factor ($Å^2$, min-max (mean)) | | | | |
| Protein | 18.86-240.60 (60.25) | 12.50-164.40 (60.74) | 54.53-210.52 (97.25) | 19.07-204.31 (61.26) |
| Nucleotides | 29.50-93.79 (55.70) | 34.94-154.19 (66.57) | 76.53-172.59 (124.10) | 29.40-152.41 (66.03) |
| Ligands | 37.90-99.40 (49.45) | 31.21-140.20 (49.15) | 63.41-98.72 (97.56) | 25.00-112.91 (68.95) |
| R.m.s deviations | | | | |
| Bond lengths (Å) | 0.003 | 0.004 | 0.002 | 0.003 |
| Bond angles (°) | 0.579 | 0.533 | 0.560 | 0.533 |
| Validation | | | | |
| MolProbity score | 1.72 | 1.59 | 1.72 | 1.61 |
| Clashscore | 9.89 | 6.94 | 6.48 | 7.63 |
| Poor rotamers (%) | 0.00 | 0.01 | 0.00 | 0.00 |
| Ramachandran plot | | | | |
| Favored (%) | 96.73 | 96.73 | 96.18 | 96.85 |
| Allowed (%) | 3.27 | 3.27 | 3.73 | 3.10 |
| Disallowed (%) | 0 | 0 | 0 | 0 |

### SUPPLEMENTARY FIGURE 1

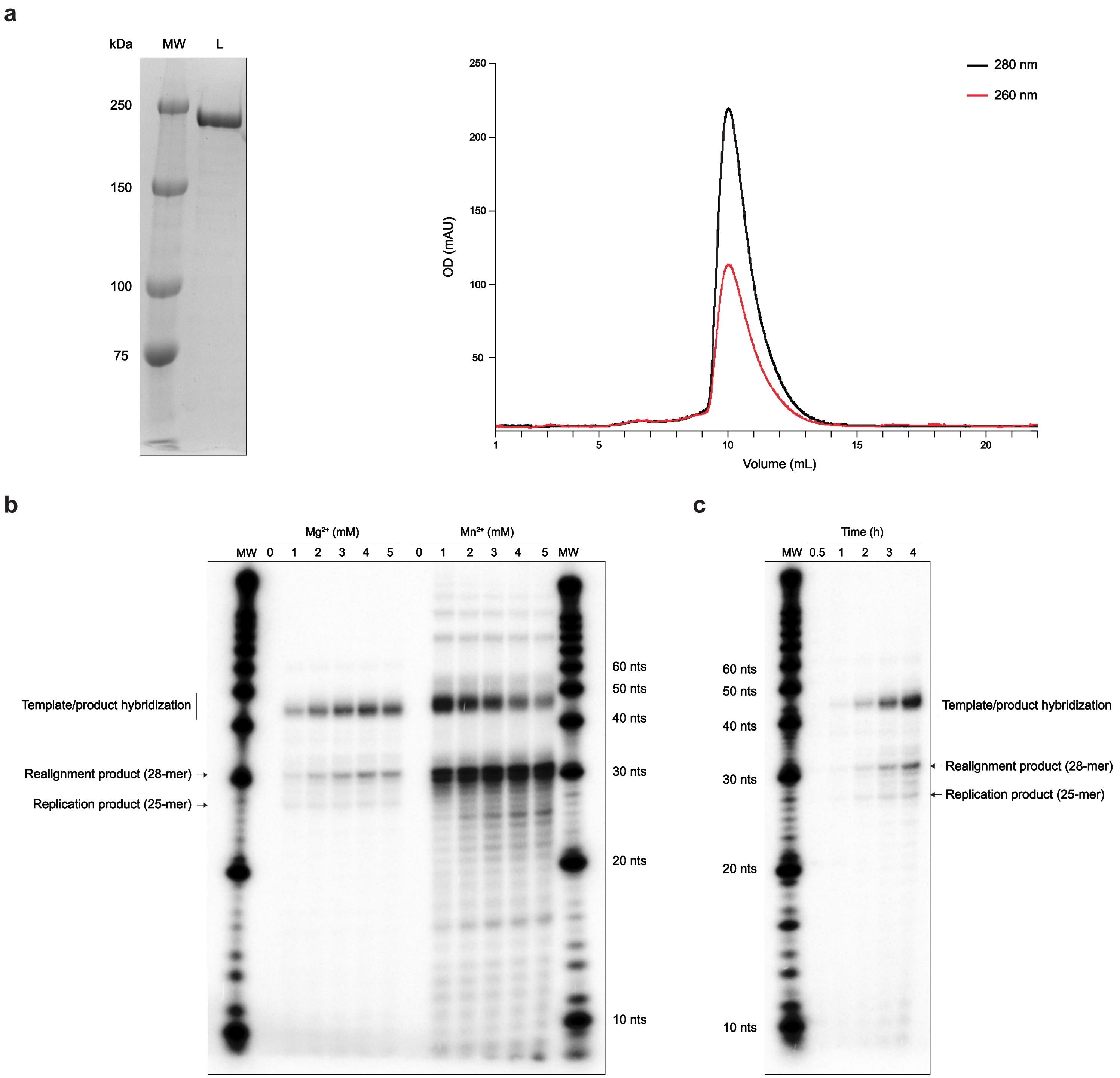


**Supplementary Fig. 1. LACV-LCItag_H34K purification and replication activity optimization**

**a**, 6% SDS-PAGE gel of LACV-LCItag_H34K (MW: Molecular weight; L: LACV-LCItag-H34K) and the corresponding gel filtration profile. Absorbance curves at 280 and 260 nm are indicated and respectively colored in black and red.

**b**, Optimization of the divalent metal ion concentration for LACV-LCItag_H34K replication activity. Assessment of replication product formation using concentration ranging from 0 to 5 mM of MgCl2 or MnCl2. The molecular weight marker (MW) corresponds to the decade marker.

**c**, Time course of LACV-LCItag_H34K replication activity. Reactions were stopped after 0.5, 1, 2, 3 or 4h.

### SUPPLEMENTARY FIGURE 2

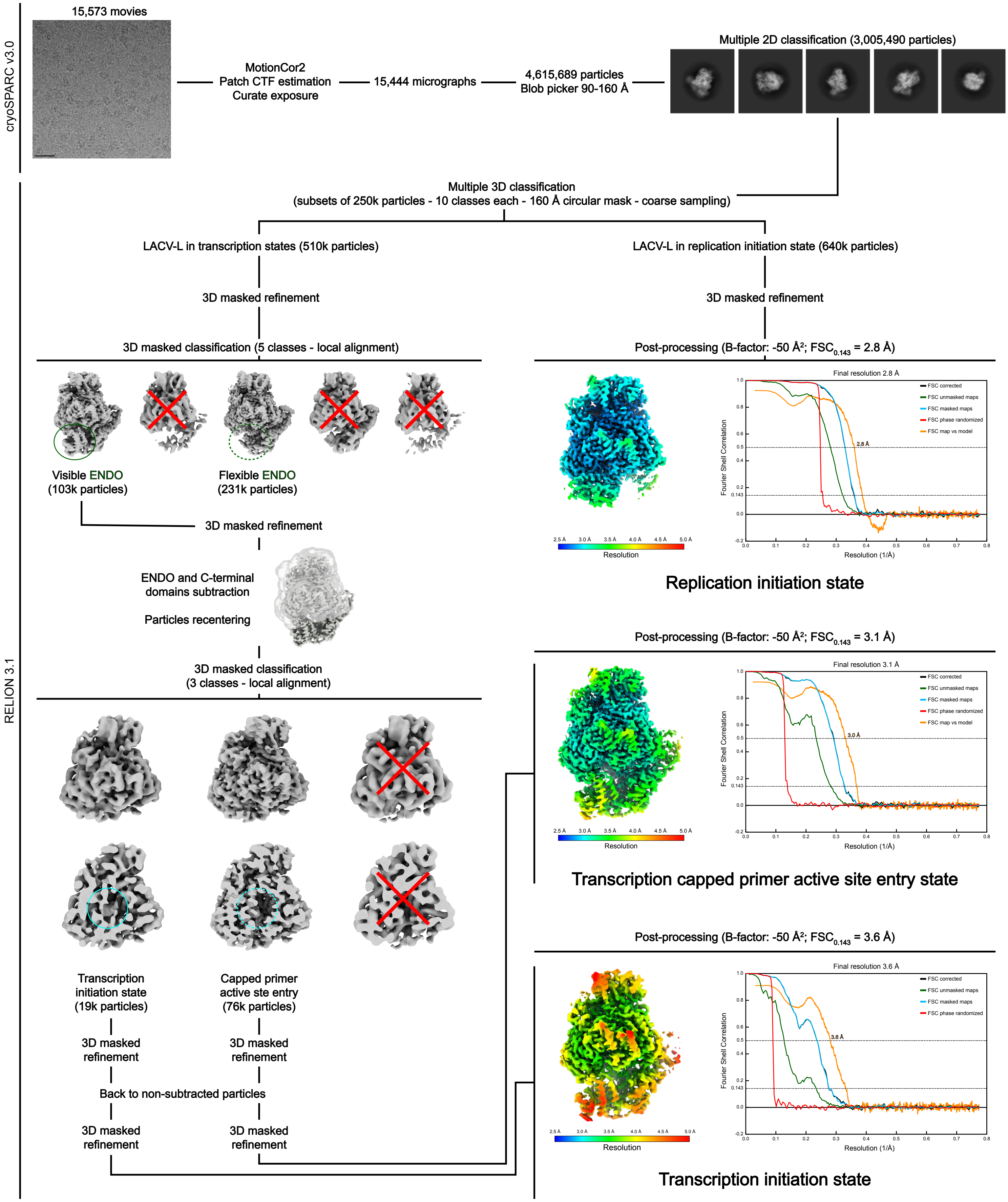


**Supplementary Fig. 2. Image processing strategy to obtain the replication initiation state, the transcription capped primer active site entry state and the transcription initiation state**

Schematics of the image processing strategy used with the data collected on a Titan Krios equipped a K3 direct electron detector. Representative micrograph, 2D class averages, 3D class averages are displayed. Local resolution EM maps colored according to resolution are shown. Fourier shell correlation curves are displayed.

### SUPPLEMENTARY FIGURE 3

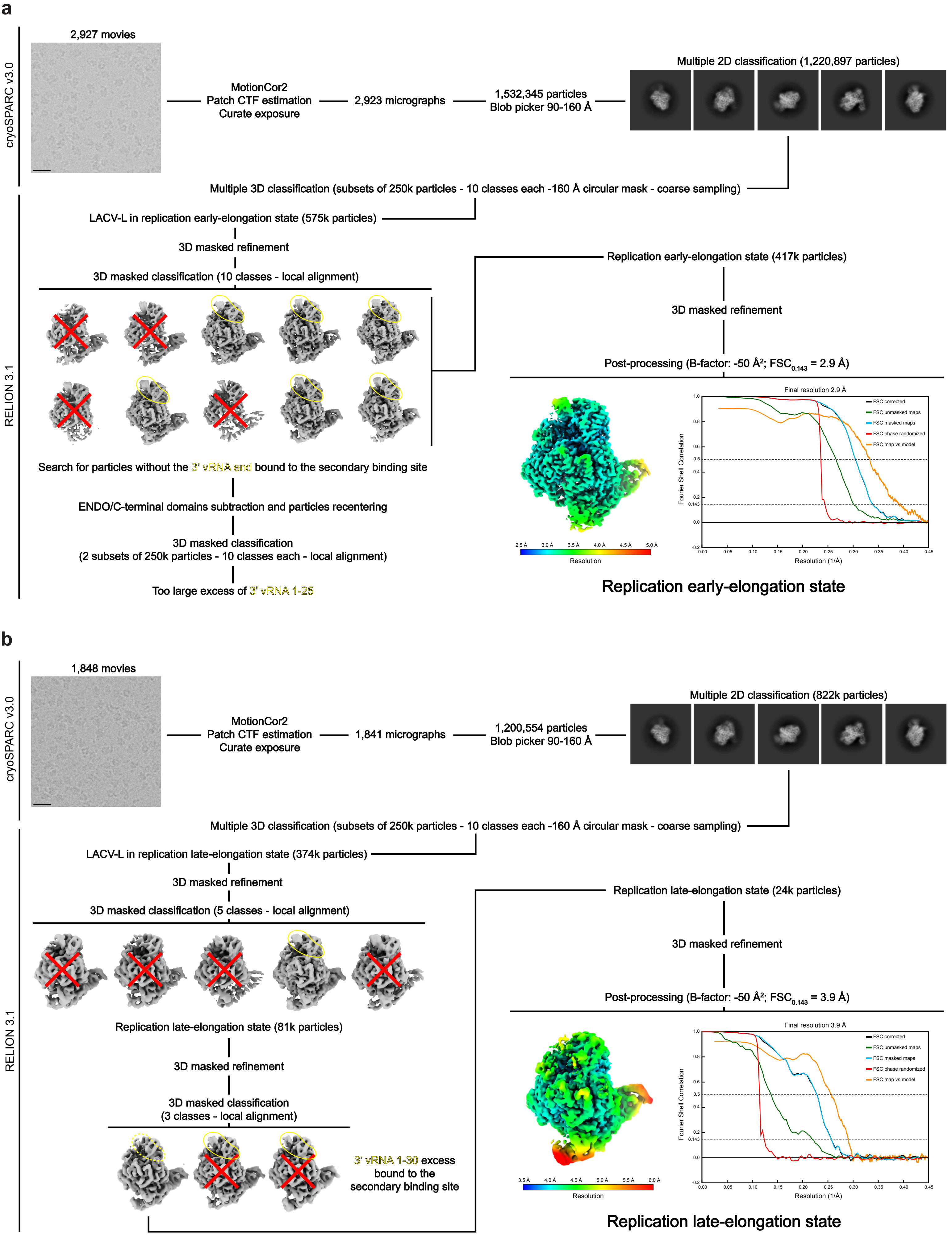


**Supplementary Fig. 3. Image processing strategy to obtain the replication early-elongation state and the replication late-elongation state**
**a**,**b**, Schematics of the image processing strategy used with the data collected on a Glacios cryo-TEM equipped a K2 direct electron detector to obtain the replication early-elongation state (a) and the replication late-elongation state (b). Representative micrographs, 2D class averages, 3D class averages are displayed. Local resolution EM maps colored according to resolution are shown. Fourier shell correlation curves are displayed.

### SUPPLEMENTARY FIGURE 4

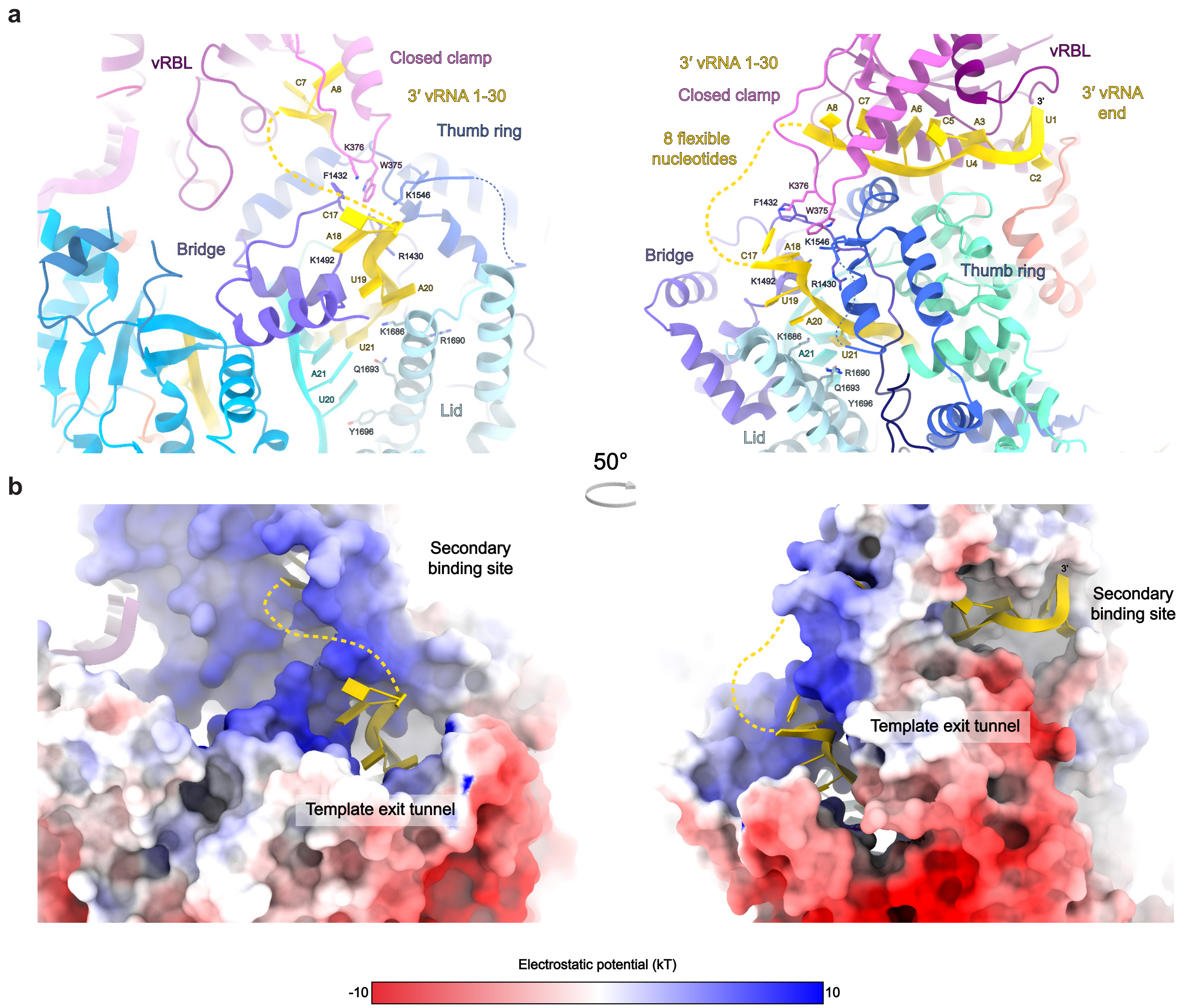


**Supplementary Fig. 4. Path of the template RNA from the template exit channel to the 3'-vRNA secondary binding site**
**a**, The 3'-vRNA is displayed in yellow. Interacting residues in the template exit channel are indicated.
**b**, LACV-LCItag_H34K surface colored according to its electrostatic potential showing that the path from the template exit channel towards the secondary binding site is positively charged.

### SUPPLEMENTARY FIGURE 5

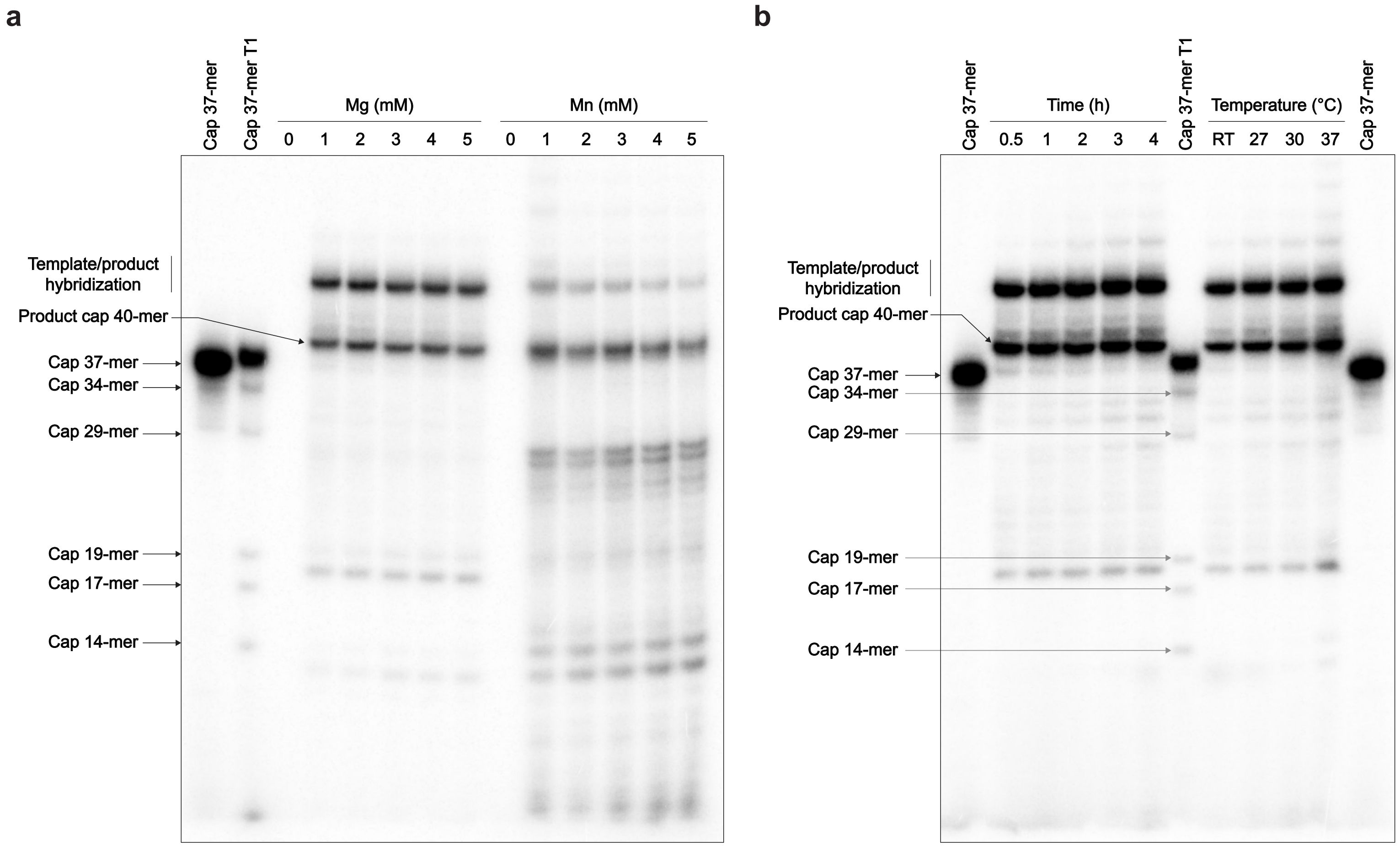


**Supplementary Fig. 5. Transcription activity optimization of LACV-LCItag_H34K**

**a**, Optimization of the divalent metal ion concentration for LACV-LCItag_H34K transcription activity. Assessment of transcription product formation using concentration ranging from 0 to 5 mM of MgCl2 or MnCl2. The Cap37-mer lane corresponds to a capped 37-mer RNA identical in sequence to the theoretical transcription product of LACV-LCItag_H34K with the cap14AG and the 3'-vRNA1-25 in the absence of realignment. The lane indicated as Cap37-mer T1 corresponds to an RNAse T1 cleavage of the cap37-mer.

**b**, Time course and temperature effect on LACV-LCItag_H34K transcription activity. Reactions were stopped after 0.5, 1, 2, 3 or 4h. Reactions were run at room temperature (RT), 27, 30 or 37°C.

### SUPPLEMENTARY FIGURE 6

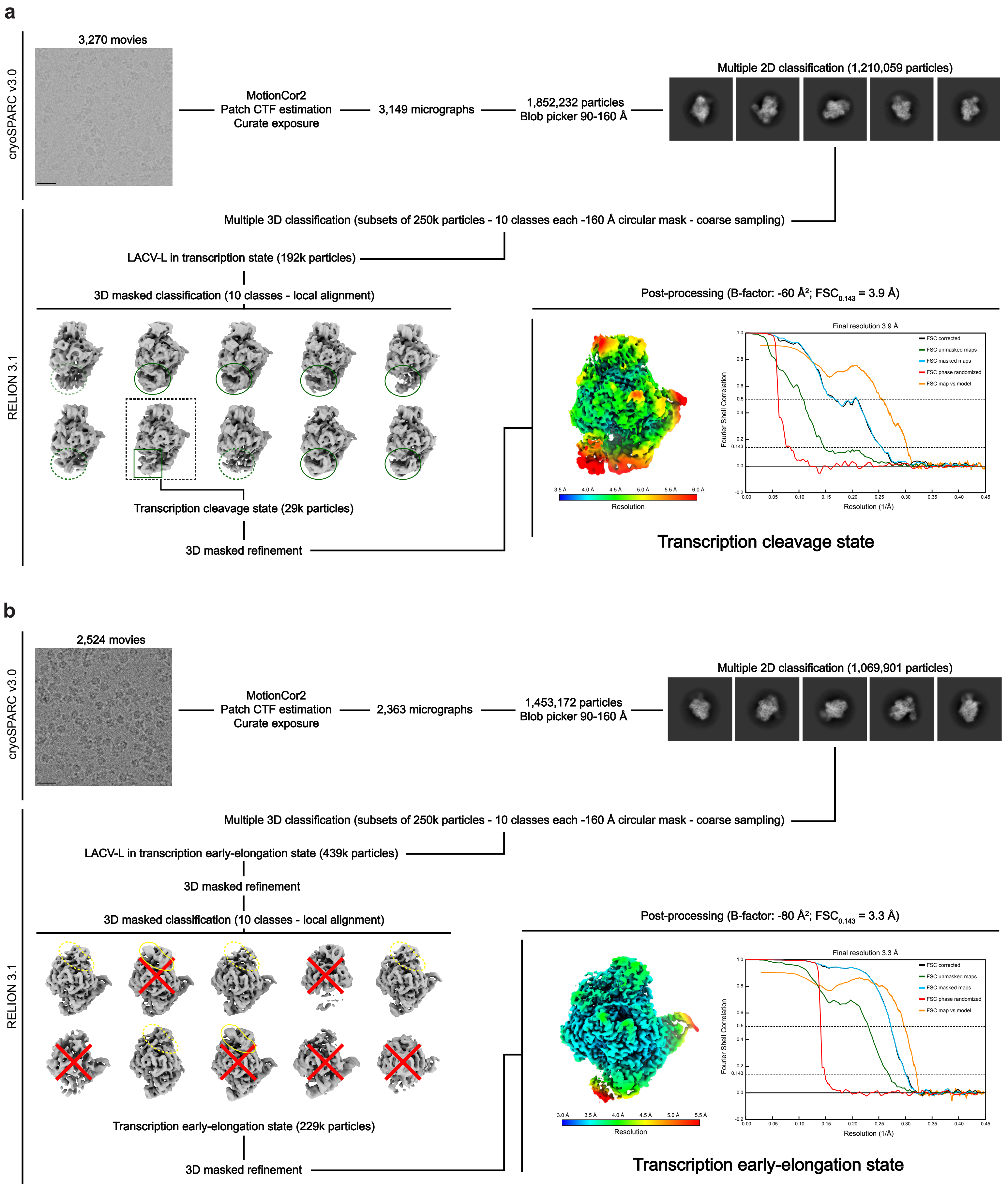


**Supplementary Fig. 6. Image processing strategy to obtain the transcription capped primer cleavage state and the transcription early-elongation state** **a**,**b**, Schematics of the image processing strategy used with the data collected on a Glacios cryo-TEM equipped a K2 direct electron detector to obtain the transcription capped primer cleavage state (a) and the transcription early-elongation state (b). Representative micrographs, 2D class averages, 3D class averages are displayed. Local resolution EM maps colored according to resolution are shown. Fourier shell correlation curves are displayed.

### SUPPLEMENTARY FIGURE 7

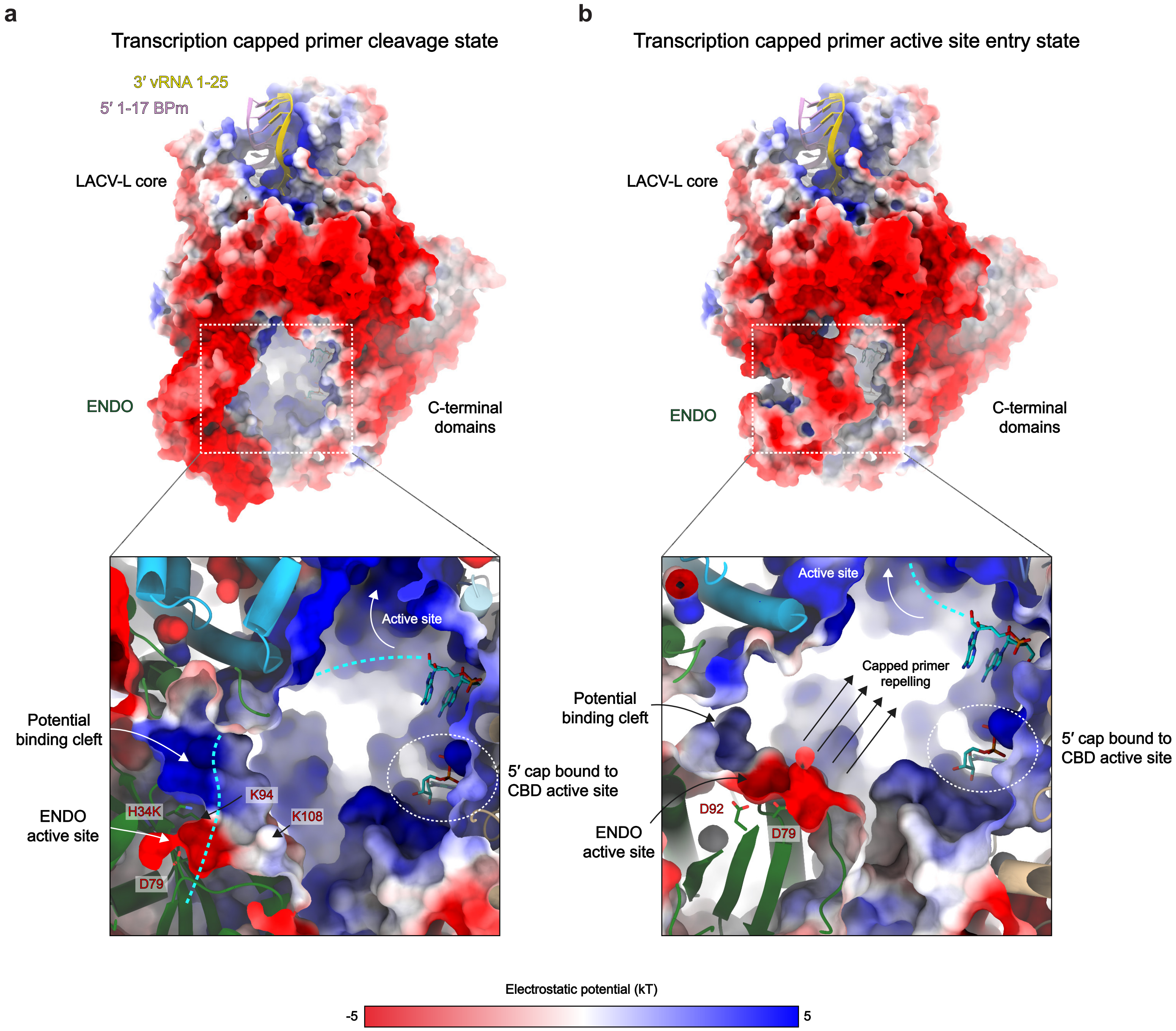


**Supplementary Fig. 7. Electrostatic surface analysis of LACV-LCltag_H34K in the transcription capped primer cleavage state and the transcription capped primer active site entry state conformations**

**a**, Electrostatic surface of the capped primer cleavage conformation. Top: global view, bottom: slabbed and zoomed view focusing on the charged environment close to the capped RNA. 5’-1-17BPm and 3’-vRNA-1-25 are displayed and respectively colored in pink and gold. RNAs tunnels are positively charged. The ENDO is presenting a potential binding cleft for the capped primer (dotted line in cyan) surrounded by K94 and K108 leading to the ENDO active site.

**b**, Electrostatic surface of the capped primer active site entry conformation as in (a). RNAs tunnels are positively charged. The ENDO active site is repelling the capped primer (dotted line in cyan) into the RdRp active site.

### SUPPLEMENTARY FIGURE 8

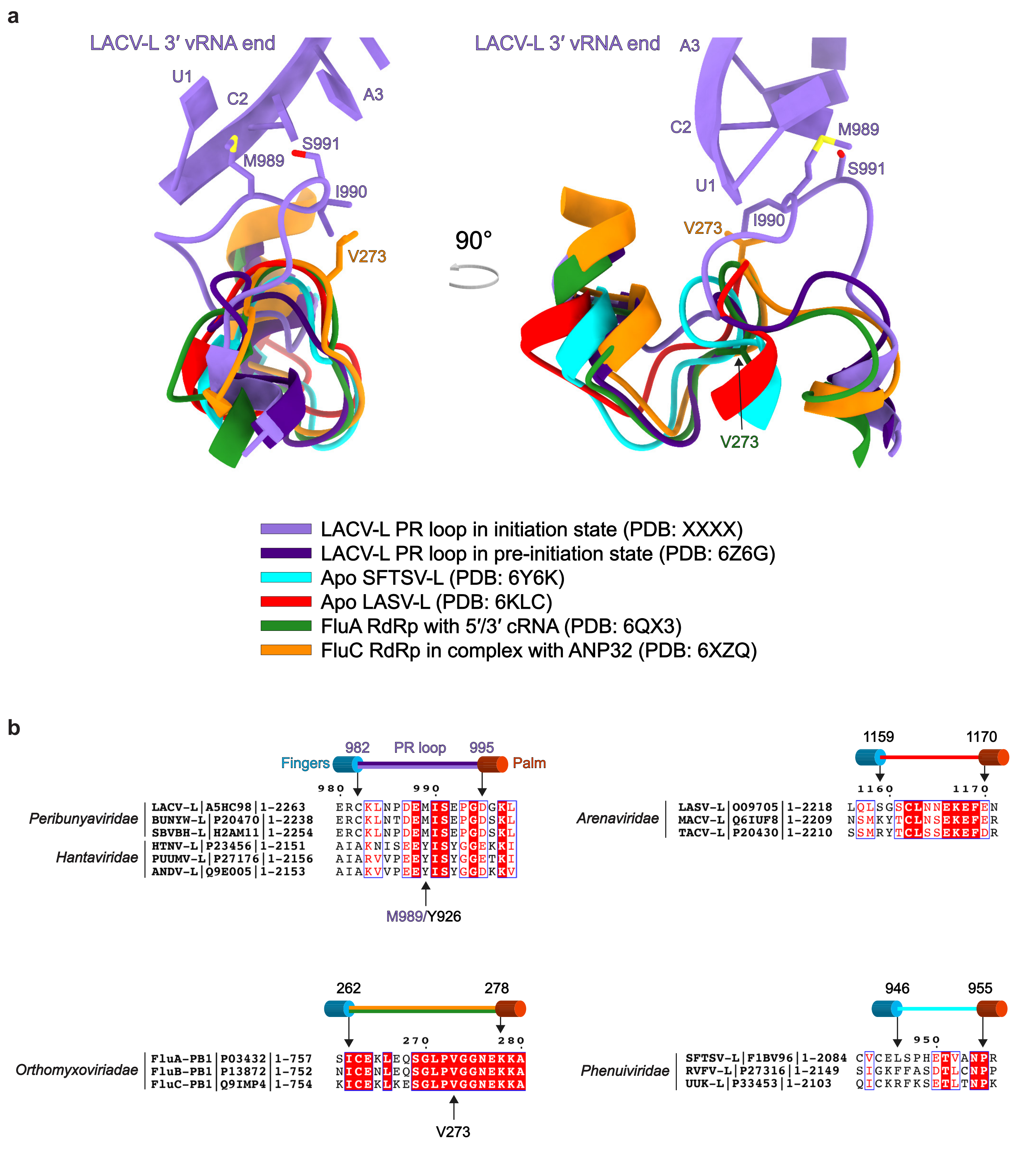


**Supplementary Fig. 8. Putative Prime-and-realign loops in sNSV**

**a**, Superposition of LACV-L PR loop in initiation state (purple), pre-initiation state (dark blue, PDB: 6Z6G), apo SFTSV-L (946-955) (light blue, PDB: 6Y6K), apo LASV-L (1159-1170) (red, PDB: 6KLC), FluA RdRp (262-278) in complex with 5'- and 3'-cRNA (green, PDB: 6QX3) and FluC RdRp (262-278) in complex with ANP32 (orange, PDB: 6XZQ). Interacting residues of the LACV-L PR loop with the 3'-vRNA end (M989, I990, S991) and the Flu RdRp (V273) are displayed.

**b**, Sequence alignment of the putative PR loop sequence in sNSV. Peribunyaviridae (LACV: La Crosse virus, BUNYW: Bunyamwera virus, SBVBH: Schmallenberg virus) and Hantaviridae (HTNV: Hantaan virus, PUUMV: Puumala virus, ANDV: Andes virus) families share some identity regarding LACV-L PR loop residues interacting with the 3'-vRNA template (E988, I990, S991). M989 in LACV-L is replaced by an aromatic residue (Y926) in Hantaviridae L protein that could play @a similar role in template stabilization. Orthomyxoviruses (Influenza A, B, C) share a conserved PR loop on PB1 subunit with V273 playing an important role in prime-and-realign mechanism (Oymans and Te Velthuis, 2018). In Arenaviridae (LASV: Lassa virus, MACV: Machupo virus, TACV: Tacaribe virus), a conserved loop is also present between residues 1159 and 1170 contrary to Phenuiviridae (SFTSV: Sever Fever with Thrombocytopenia Syndrome virus, UUK: Uukuniemi virus) in which only T951 is conserved. For each displayed sequence, the virus name, the UniProt number and the viral polymerase length are indicated.

### SUPPLEMENTARY FIGURE 9

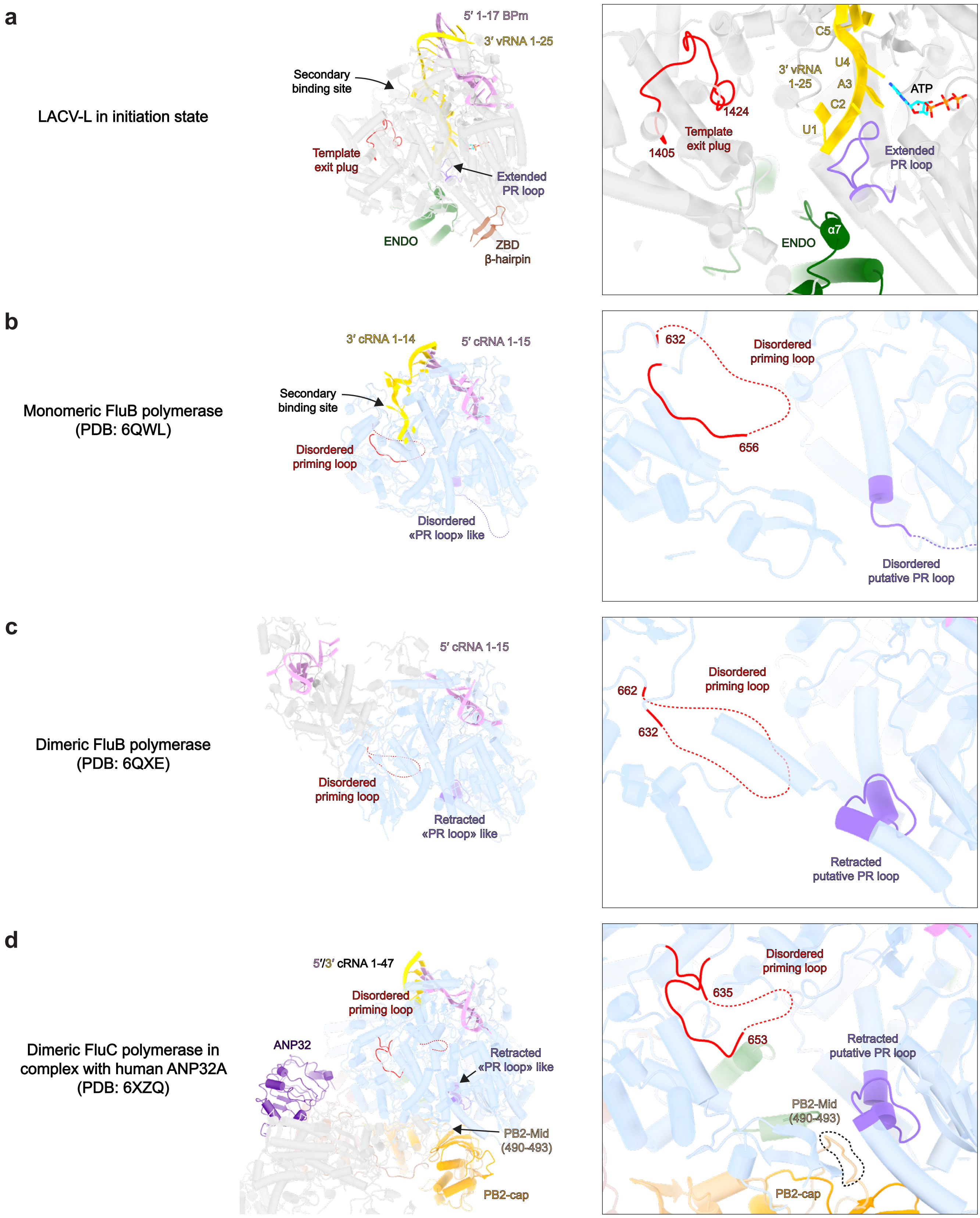


**Supplementary Fig. 9. Prime-and-realign loops, LACV-L template exit plug and influenza polymerase priming loop**
**a-d**, Cartoon representation of LACV-L in initiation state (a), monomeric influenza B polymerase bound to cRNA (PDB: 6QWL) (b), dimeric influenza B polymerase bound to cRNA (PDB: 6QXE) (c), dimeric influenza C polymerase in complex with human ANP32 (PDB: 6XZQ) (d). LACV-L core is transparent and colored in grey. Flu RdRp core is transparent and colored in blue. 3'-vRNA/cRNA end and 5'-vRNA/cRNA end present in each structure are respectively colored in gold and pink. The LACV-L template exit plug and the influenza priming loop are colored in red, if disordered represented as a dotted line. PR loops are colored in purple. Influenza PB2-cap is colored in orange, the PB2-mid in beige, both the PB2-627 and the LACV-L ZBD β-hairpin in brown. On the right panels, close-up view of the polymerase internal cavity.

### SUPPLEMENTARY FIGURE 10

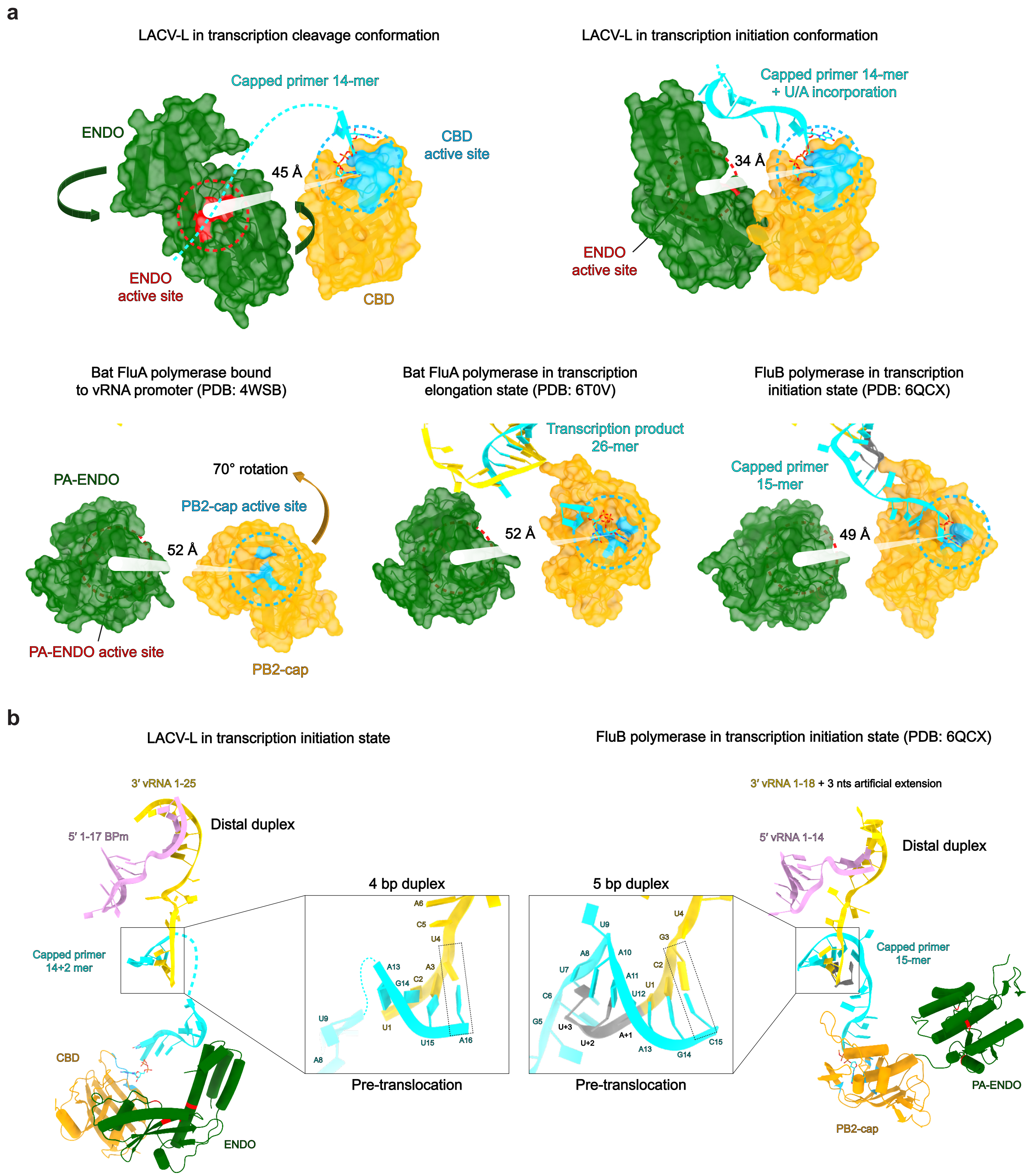


**Supplementary Fig. 10. ENDO and CBD positions in LACV-L and influenza polymerase during transcription initiation**

**a**, Surface representation of the ENDO and the CBD of LACV-L (top) and influenza polymerase (bottom). ENDOs are colored in green with active sites in red. CBDs are colored in yellow with cap-binding active sites in cyan. Distances between ENDO active sites and cap-binding active sites are indicated. The capped primer is shown in cyan and its possible path as a dotted line. Rotations between the cleavage conformation and the transcription initiation conformation are indicated with arrows.

**b**, Comparison of the paths taken by the 3'-vRNA and the capped RNA primers between LACV-L (left) and influenza B polymerase (right). Both CBD and ENDO position are displayed. A zoom is made on the nascent template-product duplexes.

#### SUPPLEMENTARY FIGURE LEGENDS

**Supplementary Fig. 1. LACV-$L_{CItag\_H34K}$ purification and replication activity optimization**

**a,** 6% SDS-PAGE gel of LACV-$L_{CItag\_H34K}$ (MW: Molecular weight; L: LACV-$L_{CItag-H34K}$) and the corresponding gel filtration profile. Absorbance curves at 280 and 260 nm are indicated and respectively colored in black and red.

**b,** Optimization of the divalent metal ion concentration for LACV-$L_{CItag\_H34K}$ replication activity. Assessment of replication product formation using concentration ranging from 0 to 5 mM of $MgCl_2$ or $MnCl_2$. The molecular weight marker (MW) corresponds to the decade marker.

**c,** Time course of LACV-$L_{CItag\_H34K}$ replication activity. Reactions were stopped after 0.5, 1, 2, 3 or 4h.

**Supplementary Fig. 2. Image processing strategy to obtain the replication initiation state, the transcription capped primer active site entry state and the transcription initiation state**

Schematics of the image processing strategy used with the data collected on a Titan Krios equipped a K3 direct electron detector. Representative micrograph, 2D class averages, 3D class averages are displayed. Local resolution EM maps colored according to resolution are shown. Fourier shell correlation curves are displayed.

**Supplementary Fig. 3. Image processing strategy to obtain the replication early-elongation state and the replication late-elongation state**

**a,b,** Schematics of the image processing strategy used with the data collected on a Glacios cryo-TEM equipped a K2 direct electron detector to obtain the replication early-elongation state **(a)** and the replication late-elongation state **(b)**. Representative micrographs, 2D class averages, 3D class averages are displayed. Local resolution EM maps colored according to resolution are shown. Fourier shell correlation curves are displayed.

**Supplementary Fig. 4. Path of the template RNA from the template exit channel to the 3'-vRNA secondary binding site**

**a,** The 3'-vRNA is displayed in yellow. Interacting residues in the template exit channel are indicated.

**b,** LACV-$L_{CItag\_H34K}$ surface colored according to its electrostatic potential showing that the path from the template exit channel towards the secondary binding site is positively charged.

**Supplementary Fig. 5. Transcription activity optimization of LACV-$L_{Cltag\_H34K}$**

**a,** Optimization of the divalent metal ion concentration for LACV-$L_{Cltag\_H34K}$ transcription activity. Assessment of transcription product formation using concentration ranging from 0 to 5 mM of $MgCl_2$ or $MnCl_2$. The Cap37-mer lane corresponds to a capped 37-mer RNA identical in sequence to the theoretical transcription product of LACV-$L_{Cltag\_H34K}$ with the cap14AG and the 3'-vRNA1-25 in the absence of realignment. The lane indicated as Cap37-mer T1 corresponds to an RNAse T1 cleavage of the cap37-mer.

**b,** Time course and temperature effect on LACV-$L_{Cltag\_H34K}$ transcription activity. Reactions were stopped after 0.5, 1, 2, 3 or 4h. Reactions were run at room temperature (RT), 27, 30 or 37°C.

**Supplementary Fig. 6. Image processing strategy to obtain the transcription capped primer cleavage state and the transcription early-elongation state**

**(a-b)** Schematics of the image processing strategy used with the data collected on a Glacios cryo-TEM equipped a K2 direct electron detector to obtain the transcription capped primer cleavage state **(a)** and the transcription early-elongation state **(b)**. Representative micrographs, 2D class averages, 3D class averages are displayed. Local resolution EM maps colored according to resolution are shown. Fourier shell correlation curves are displayed.

**Supplementary Fig. 7. Electrostatic surface analysis of LACV-$L_{Cltag\_H34K}$ in the transcription capped primer cleavage state and the transcription capped primer active site entry state conformations**

**a,** Electrostatic surface of the capped primer cleavage conformation. Top: global view, bottom: slabbed and zoomed view focusing on the charged environment close to the capped RNA. 5'-1-17BPm and 3'-vRNA-1-25 are displayed and respectively colored in pink and gold. RNAs tunnels are positively charged. The ENDO is presenting a potential binding cleft for the capped primer (dotted line in cyan) surrounded by K94 and K108 leading to the ENDO active site.

**b,** Electrostatic surface of the capped primer active site entry conformation as in **(a)**. RNAs tunnels are positively charged. The ENDO active site is repelling the capped primer (dotted line in cyan) into the RdRp active site.

**Supplementary Fig. 8. Putative Prime-and-realign loops in sNSV**

**a,** Superposition of LACV-L PR loop in initiation state (purple), pre-initiation state (dark blue, PDB: 6Z6G), apo SFTSV-L (946-955) (light blue, PDB: 6Y6K), apo LASV-L (1159-1170) (red, PDB: 6KLC), FluA RdRp (262-278) in complex with 5'- and 3'-cRNA (green, PDB: 6QX3) and FluC RdRp (262-278) in complex with ANP32 (orange, PDB: 6XZQ). Interacting residues of the LACV-L PR loop with the 3'-vRNA end (M989, I990, S991) and the Flu RdRp (V273) are displayed.

**b,** Sequence alignment of the putative PR loop sequence in sNSV. *Peribunyaviridae* (LACV: La Crosse virus, BUNYW: Bunyamwera virus, SBVBH: Schmallenberg virus) and *Hantaviridae* (HTNV: Hantaan virus, PUUMV: Puumala virus, ANDV: Andes virus) families share some identity regarding LACV-L PR loop residues interacting with the 3'-vRNA template (E988, I990, S991). M989 in LACV-L is replaced by an aromatic residue (Y926) in *Hantaviridae* L protein that could play a similar role in template stabilization. Orthomyxoviruses (Influenza A, B, C) share a conserved PR loop on PB1 subunit with V273 playing an important role in prime-and-realign mechanism (Oymans and Te Velthuis, 2018). In *Arenaviridae* (LASV: Lassa virus, MACV: Machupo virus, TACV: Tacaribe virus), a conserved loop is also present between residues 1159 and 1170 contrary to *Phenuiviridae* (SFTSV: Sever Fever with Thrombocytopenia Syndrome virus, UUK: Uukuniemi virus) in which only T951 is conserved. For each displayed sequence, the virus name, the UniProt number and the viral polymerase length are indicated.

**Supplementary Fig. 9. Prime-and-realign loops, LACV-L template exit plug and influenza polymerase priming loop**

**a-d,** Cartoon representation of LACV-L in initiation state **(a)**, monomeric influenza B polymerase bound to cRNA (PDB: 6QWL) **(b)**, dimeric influenza B polymerase bound to cRNA (PDB: 6QXE) **(c)**, dimeric influenza C polymerase in complex with human ANP32 (PDB: 6XZQ) **(d)**. LACV-L core is transparent and colored in grey. Flu RdRp core is transparent and colored in blue. 3'-vRNA/cRNA end and 5'-vRNA/cRNA end present in each structure are respectively colored in gold and pink. The LACV-L template exit plug and the influenza priming loop are colored in red, if disordered represented as a dotted line. PR loops are colored in purple. Influenza PB2-cap is colored in orange, the PB2-mid in beige, both the PB2-627 and the LACV-L ZBD β-hairpin in brown. On the right panels, close-up view of the polymerase internal cavity.

**Supplementary Fig. 10. ENDO and CBD positions in LACV-L and influenza polymerase during transcription initiation**

**a,** Surface representation of the ENDO and the CBD of LACV-L (top) and influenza polymerase (bottom). ENDOs are colored in green with active sites in red. CBDs are colored in yellow with cap-binding active sites in cyan. Distances between ENDO active sites and cap-binding active sites are indicated. The capped primer is shown in cyan and its possible path as a dotted line. Rotations between the cleavage conformation and the transcription initiation conformation are indicated with arrows.

**b,** Comparison of the paths taken by the 3'-vRNA and the capped RNA primers between LACV-L (left) and influenza B polymerase (right). Both CBD and ENDO position are displayed. A zoom is made on the nascent template-product duplexes.

**Supplementary Movie 1. Replication: LACV-L from pre-initiation to late-elongation**

**Supplementary Movie 2. Transcription: LACV-L from pre-initiation to early-elongation**
